# Analysis of single-cell RNA sequencing data based on autoencoders

**DOI:** 10.1101/727867

**Authors:** Andrea Tangherloni, Federico Ricciuti, Daniela Besozzi, Pietro Liò, Ana Cvejic

**Affiliations:** Wellcome Trust-Medical Research Council Cambridge Stem Cell Institute, Cambridge, UK; Department of Haematology, University of Cambridge, Cambridge, UK; Wellcome Trust Sanger Institute, Wellcome Trust Genome Campus, Hinxton, UK; Department of Human and Social Sciences, University of Bergamo, Bergamo, Italy; Department of Informatics, Systems and Communication, University of Milano-Bicocca, Milan, Italy; Bicocca Bioinformatics, Biostatistics and Bioimaging Centre (B4), Milan, Italy; Department of Computer Science and Technology, University of Cambridge, Cambridge, UK

**Keywords:** Autoencoders, scRNA-Seq, Dimensionality reduction, Clustering, Batch correction, Data integration

## Abstract

**Background:** Single-cell RNA sequencing (scRNA-Seq) experiments are gaining ground to study the molecular processes that drive normal development as well as the onset of different pathologies. Finding an effective and efficient low-dimensional representation of the data is one of the most important steps in the downstream analysis of scRNA-Seq data, as it could provide a better identification of known or putatively novel cell-types. Another step that still poses a challenge is the integration of different scRNA-Seq datasets. Though standard computational pipelines to gain knowledge from scRNA-Seq data exist, a further improvement could be achieved by means of machine learning approaches.

**Results:** Autoencoders (AEs) have been effectively used to capture the non-linearities among gene interactions of scRNA-Seq data, so that the deployment of AE-based tools might represent the way forward in this context. We introduce here scAEspy, a unifying tool that embodies: (1) four of the most advanced AEs, (2) two novel AEs that we developed on purpose, (3) different loss functions. We show that scAEspy can be coupled with various batch-effect removal tools to integrate data by different scRNA-Seq platforms, in order to better identify the cell-types. We benchmarked scAEspy against the most used batch-effect removal tools, showing that our AE-based strategies outperform the existing solutions.

**Conclusions:** scAEspy is a user-friendly tool that enables using the most recent and promising AEs to analyse scRNA-Seq data by only setting up two user-defined parameters. Thanks to its modularity, scAEspy can be easily extended to accommodate new AEs to further improve the downstream analysis of scRNA-Seq data. Considering the relevant results we achieved, scAEspy can be considered as a starting point to build a more comprehensive toolkit designed to integrate multi single-cell omics.

## Background

Single-cell RNA sequencing (scRNA-Seq) was named the “Method of the Year” in 2013, and it is currently used to investigate cell-to-cell heterogeneity since it allows for measuring the transcriptome-wide gene expression at single-cell resolution, enabling the identification of different cell-types. scRNA-Seq data are prevalent generated in studies that aim at understanding the molecular processes driving normal development and the onset of pathologies [1, 2]. This field of research continuously poses new computational questions that have to be addressed [3].

One of the most important steps in scRNA-Seq analysis is the clustering of cells into groups that correspond to known or putatively novel cell-types, by considering the expression of common sets of signature genes. However, this step still remains a challenging task because applying clustering approaches in high-dimensional spaces can generate misleading results, as the distance between most pairs of points is similar [4]. As a consequence, finding an effective and efficient low-dimensional representation of the data is one of the most crucial steps in the downstream analysis of scRNA-Seq data. A common workflow of downstream analysis, depicted in Figure 1, includes two dimensionality reduction steps: (1) Principal Component Analysis (PCA) [5] for an initial reduction of the dimensions based on the Highly Variable Genes (HVGs), and (2) a non-linear dimensionality reduction approach— e.g., t-distributed Stochastic Neighbour Embedding (t-SNE) [6] or Uniform Manifold Approximation and Projection (UMAP) [7]—on the PCA space for visualisation purposes (e.g., showing the labelled clusters) [8, 9]. In addition, when multiple scRNA-Seq datasets have to be combined for further analyses, the technical non-negligible batch-effects that may exist among the datasets must be taken into account [3, 8, 10–13], making the dimensionality reduction even more complicated and fundamental. Indeed, finding a salient batch corrected and a low dimensional embedding space can help to better partition and distinguish the various cell-types. Although commonly used approaches for dimensionality reduction achieved good performance when applied to scRNA-Seq data [8], novel and more robust dimensionality reduction strategies should be used to account for the sparsity, intrinsic noise, unexpected dropout, and burst effects [3, 14], as well as the low amounts of RNA that are typically present in single-cells. Ding *et al*. showed that low-dimensional representations of the original data learned using latent variable models preserve both the local and global neighbour structures of the original data [15]. Autoencoders (AEs) showed outstanding performance in this regard due to their ability to capture the strong non-linearities among the gene interactions existing in the high-dimensional expression space.

**Figure 1.**
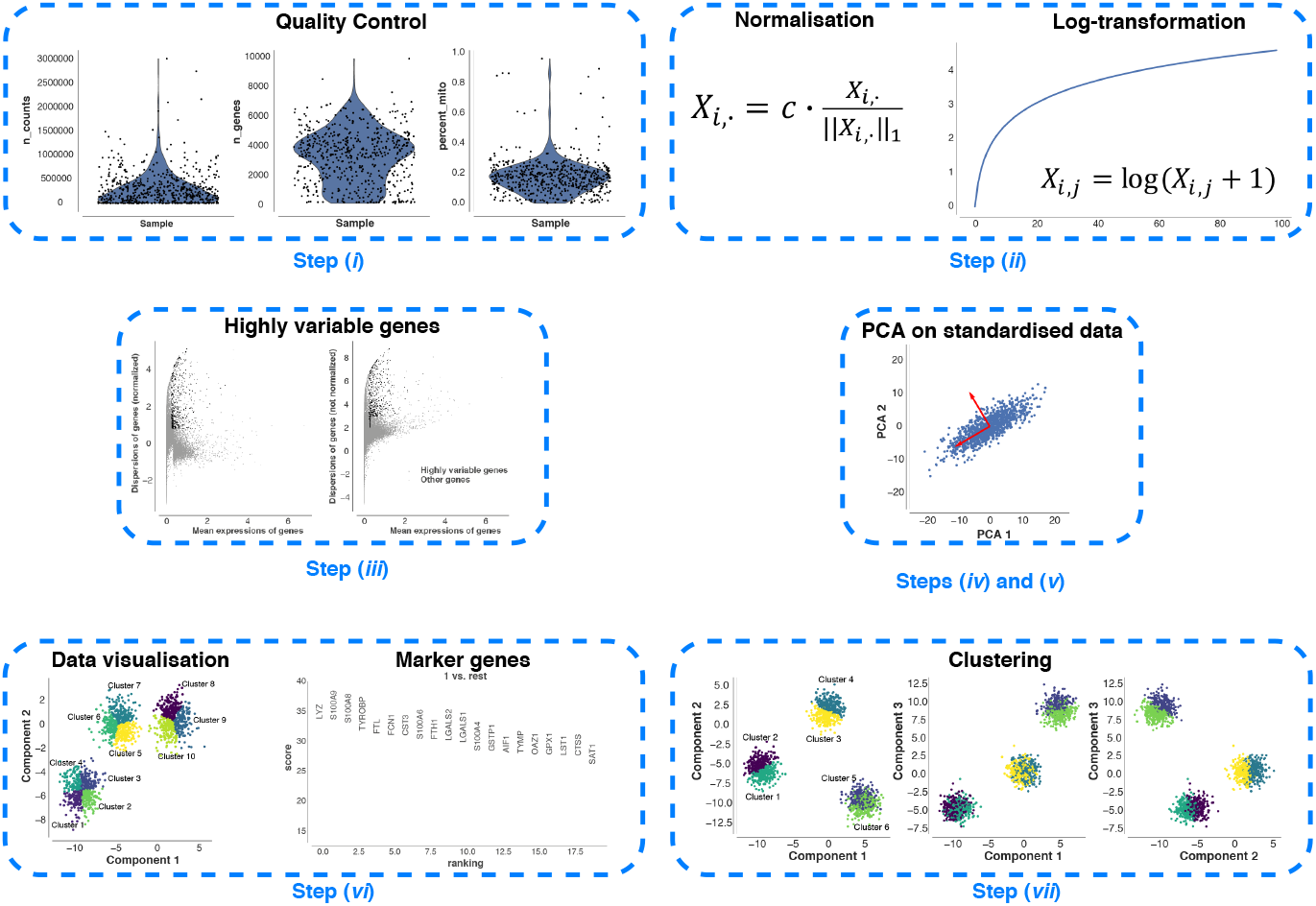
A common workflow for the downstream analysis of scRNA-Seq data. The workflow includes the following seven steps: (*i*) quality control to remove low-quality cells that may add technical noise, which could obscure the real biological signals; (*ii*) normalisation and log-transformation; (*iii*) identification of the HVGs to reduce the dimensionality of the dataset by including only the most informative genes; (*iv*) standardisation of each gene to zero mean and unit variance; (*v*) dimensionality reduction generally obtained by applying PCA; (*vi*) clustering of the cells starting from the low-dimensional representation of the data that are used to annotate the obtained clusters (i.e., identification of known and putatively novel cell-types); (*vii*) data visualisation on the low-dimensional space generated by applying a non-linear approach (e.g., SNE or UMAP) on the reduced space calculated in step (*v*).

### Autoencoders for denoising and dimensionality reduction

Deep Count AE network (DCA) was one of the first AE-based approach proposed to denoise scRNA-Seq datasets [16] by considering the count distribution, overdispersion, and sparsity of the data. DCA relies on a negative binomial noise model, with or without zero-inflation, to capture non-linear gene-gene dependencies. Starting from the vanilla version of the Variational AE (VAE) [17], several approaches have been proposed. Among them, single-cell Variational Inference (scVI) was the first scalable framework that allowed for a probabilistic representation and analysis of gene expression datasets [18]. scVI was built upon Deep Neural Networks (DNNs) and stochastic optimization to consider the information across similar cells and genes to approximate the distributions underlying the analysed gene expression data. This computational tool allows for coupling low-dimensional probabilistic representation of gene expression data with the downstream analysis to consider the measurement of uncertainty through a statistical model. Svensson *et al*. integrated a Linearly Decoded VAE (LDVAE) into scVI [19], enabling the identification of relationships among the cell representation coordinates and gene weights via a factor mode.

Single-cell VAE (scVAE) was introduced to directly model the raw counts from RNA-seq data [20]. More importantly, the authors proposed a Gaussian-mixture model to better learn biologically plausible groupings of scRNA-Seq data on the latent space.

The framework called Decomposition using Hierarchical AE (scDHA) is composed of two modules [21]. The first module is a non-negative kernel AE able to provide a non-negative, part-based denoised representation of the original data. During this step, the genes and the components having an insignificant contribution to the denoised representation of the data are removed. The second module is a stacked Bayesian self-learning network built upon the VAE. This specific module is used to project the denoised data into a low-dimensional space used during the downstream analysis. scDHA outperformed PCA, t-SNE, and UMAP in terms of silhouette index [22] on the tested datasets.

AEs coupled with disentanglement methods have been used to both improve the data representation and obtain better separation of the biological factors of variation in gene expression data [23]. In addition, a graph AE, consisting of graph convolutional layers, was developed to predict relationships between single-cells. This framework can be used to identify the cell-types in the dataset under analysis and discover the driver genes for the differentiation process. Wang *et al*. proposed a deep VAE for scRNA-Seq data named VASC [24], a deep multi-layer generative model that improves the dimensionality reduction and visualisation steps in an unsupervised manner. Thanks to its ability to model dropout events—that can hinder various downstream analysis steps (e.g., clustering analysis, differential expression analysis, inference of gene-to-gene relationships) by introducing a high number of zero counts in the expression matrices—and to find non-linear hierarchical representations of the data, VASC obtained superior performance with respect to four state-of-the-art dimensionality reduction and visualisation approaches [24].

Dimensionality Reduction with Adversarial VAE (DR-A) has been recently proposed to fulfil the dimensionality reduction step from a data-driven point of view [25]. Compared to the previous approaches, DR-A exploits an adversarial VAE-based framework, which is a recent variant of generative adversarial networks. DR-A generally obtained more accurate low-dimensional representation of scRNA-Seq data compared to state-of-the-art approaches (e.g., PCA, scVI, t-SNE, UMAP), leading to better clustering performance. Geddes *et al*. proposed an AE-based cluster ensemble framework to improve the clustering process [26]. As a first step, random subspace projections of the data are compressed onto a low-dimensional space by exploiting an AE, obtaining different encoded spaces. Then, an ensemble clustering approach is applied across all the encoded spaces to generate a more accurate clustering of the cells.

### Autoencoders for the imputation of missing data

AutoImpute was proposed to deal with the insufficient quantities of starting RNA in the individual cells, a problem that generally leads to significant dropout events. As a consequence, the resulting gene expression matrices are sparse and contain a high number of zero counts. AutoImpute is an AE-based imputation method that works on sparse gene expression matrices, trying to learn the inherent distribution of the input data to assign the missing values [27]. scSVA was also proposed to identify and recover dropout events [28], which are imputed by fitting a mixed model of each possible cell-type. In addition, it performs an efficient feature extraction step of the high-dimensional scRNA-Seq data, obtaining a low-dimensional embedding. In the tests showed by the authors, scSVA was able to outperform different state-of-the-art and novel approaches (e.g., PCA, t-SNE, UMAP, VASC).

Other two methods based on non-parametric AEs were proposed to address the imputation problem [29]. Learning with AuToEncoder (LATE) relies on an AE that is directly trained on a gene expression matrix with parameters randomly generated, while TRANSfer learning with LATE (TRANSLATE) takes into consideration a reference gene expression dataset to estimate the parameters that are then used by LATE on the new gene expression matrix. LATE and TRANSLATE were able to obtain outstanding performance on both real and simulated data by recovering non-linear relationships in pairs of genes, allowing for a better identification and separation of the cell-types.

GraphSCI combines Graph convolution network and AE,it is the first approach that systematically integrates gene-to-gene relationships with the gene expression data into a deep learning framework. GraphSCI is able to impute the dropout events by taking advantage of low-dimensional representations of similar cells and gene-gene interactions [30].

Generally, in the existing AEs the input data are usually codified in a specific format, making their integration into the existing scRNA-Seq analysis toolkits (e.g., Scanpy [31] and Seurat [32]) a difficult task. In addition, the existing tools are implemented in Keras (https://github.com/fchollet/keras), TensorFlow [33] or PyTorch [34], and all the three libraries are thus required to run them. Finally, the currently available AEs cannot be directly exploited to obtain the latent space or to generate synthetic cells. In order to overcome the described limitations, we developed scAEspy, which is a unifying, user-friendly, and standalone tool that relies only on TensorFlow and allows easy access to different AEs by setting up only two user-defined parameters. scAEspy can be used on High-Performance Computing (HPC) infrastructures to speed-up its execution. It can be easily run on clusters of both Central Processing Units (CPUs) and Graphics Processing Units (GPUs). Indeed, it was designed and developed to be executed on multi- and many-core infrastructures. In addition, scAEspy gives access to the latent space, generated by the trained AE, which can be directly used to show the cells in this embedded space or as a starting point for other dimensionality reduction approaches (e.g., t-SNE and UMAP) as well as downstream analyses (e.g., batch-effect removal).

In this work, we show how scAEspy can be used to deal with the existing batch-effects among samples. Indeed, the application of batch-effect removal tools into the latent space allowed us to outperform state-of-the-art methods as well as the same batch-effect removal tools applied on the PCA space. Finally, scAEspy implements different loss functions, which are fundamental to deal with different sequencing platforms.

## Results

We tested PCA and AEs to address the integration of different datasets. Specifically, we used all the AEs implemented in our scAEspy tool: VAE [17], an AE only based on the Maximum Mean Discrepancy (MMD) distance (called here MMDAE) [35], MMDVAE, Gaussian-mixture VAE (GMVAE), and two novel Gaussian-mixture AEs that we developed, called GMMMD and GMMMDVAE, respectively. In all the performed tests, the constrained versions of the following loss functions were used: Negative Binomial (NB), Poisson, zero-inflated NB (ZINB), zero-inflated Poisson (ZIP). We used a number of Gaussian distributions equal to the number of datasets to integrate for GMVAE, GMMMD, and GMMMDVAE. In addition, we tested the following configurations of hidden layer and latent space to understand how the dimension of the AEs might potentially affect the performance: (256, 64), (256, 32), (256, 16), (128, 64), (128, 32), (128, 16), (64, 32), and (64, 16), where (*H, L*) represents the number of neurons composing the hidden layer (*H* neurons) and latent space (*L* neurons).

In order to deal with the possible batch-effects, we applied the following approaches, as suggested in [8,11] and being the most used batch-effect removal tools in the literature: Batch Balanced *k*-Nearest Neighbours (BBKNN) [36], Harmony [37], ComBat [38–40], and the Seurat implementation of the Canonical Correlation Analysis (CCA) [13]. Thus, we compared vanilla PCA and AEs, PCA and AEs followed by either BBKNN or Harmony, ComBat, and CCA. For each batch-effect removal tool, we used different samples that have to be merged as batches.

The proposed strategies were compared on three publicly available datasets, namely: Peripheral Blood Mononuclear Cells (PBMCs), Pancreatic Islet Cells (PICs), and Mouse Cell Atlas (MCA) by using well-known clustering metrics (i.e., Adjusted Rand Index, Adjusted Mutual Information Index, Fowlkes Mallows Index, Homogeneity Score, and V-Measure). It is worth mentioning that generally the cell-types are manually identified by expert biologists starting from an overclustering or underclustering of the data, possibly followed by different steps of subclustering of some clusters. Here, we evaluate how the different strategies are able to automatically separate the cells by fixing the number of clusters equal to the number of cell-types manually identified by the authors of the papers.

## Datasets

### Peripheral Blood Mononuclear Cells

PBMCs from eight patients with systemic lupus erythematosus were collected and processed using the 10× Chromium Genomics platform [41]. The dataset is composed of a control group (6573 cells) and an interferon-*β* stimulated group (7466 cells). We considered the 8 distinct cell-types identified by the authors following a standard workflow [41]. The count matrices were downloaded from Seurat’s tutorial *“Integrating stimulated vs. control PBMC datasets to learn cell-type specific responses”* (https://satijalab.org/seurat/v3.0/immune_alignment.html).

### Pancreatic islet cells

PIC datasets were generated independently using four different platforms: CEL-Seq [42] (1004 cells), CEL-Seq2 [43] (2285 cells), Fluidigm C1 [44] (638 cells), and Smart-Seq2 [45] (2394 cells). For our tests, we considered the 13 different cell-types across the datasets identified in [46] by applying PCA on the scaled integrated data matrix. The count matrices were downloaded from Seurat’s tutorial *“Integration and Label Transfer”* (https://satijalab.org/seurat/v3.0/integration.html).

### Mouse cell atlas

MCA is composed of two different datasets. The former was generated by Han *et al*. [47] using Microwell-Seq (4239 cells) [47], while the latter by the Tabula Muris Consortium [48] using Smart-Seq2 (2715 cells). The 11 distinct cell-types with the highest number of cells, which were present in both datasets, have been taken into account as in [11]. The count matrices were downloaded from the public GitHub repository related to [11] (https://github.com/JinmiaoChenLab/Batch-effect-removal-benchmarking).

### Metrics

#### Adjusted Rand Index

The Rand Index (RI) is a similarity measure between the results obtained from the application of two different clustering methods. The first clustering method is used as ground truth (i.e., true clusters), while the second one has to be evaluated (i.e., predicted clusters). The RI is calculated by considering all pairs of samples appearing in the clusters, namely, it counts the pairs that are assigned either to the same or different clusters in both the predicted and the true clusters. The Adjusted RI (ARI) [49] is the “adjusted for chance” version of the RI. Its values vary in the range [*−*1, 1]: a value close to 0 means a random assignment, independently of the number of clusters, while a value equal to 1 indicates that the clusters obtained with both clustering approaches are identical. Negative values are obtained if the index is less than the expected index.

#### Adjusted Mutual Information Index

The Mutual Information Index (MII) [50] represents the mutual information of two random variables, which is a similarity measure of the mutual dependence between the two variables. Specifically, it is used to quantify the amount of information that can be gained by one random variable observing the other variable. The MII is strictly correlated with the entropy of a random variable, which quantifies the expected “amount of information” that is contained in a random variable. This index is used to measure the similarity between two labels of the same data. Similarly to ARI, the Adjusted MII (AMII) is “adjusted for chance” and its values vary in the range [0, 1].

#### Fowlkes Mallows Index

The Fowlkes Mallows Index (FMI) [51] measures the similarity between the clusters obtained by using two different clustering approaches. It is defined as the geometric mean between precision and recall. Assuming that the first clustering approach is the ground truth, the precision is the percentage of the results that are relevant, while the recall refers to the percentage of total relevant results correctly assigned by the second clustering approach. The index ranges from 0 to 1.

#### Homogeneity Score

The result of the tested clustering approach satisfies the Homogeneity Score (HS) [52] if all of its clusters contain only cells that belong to a single cell-type. Its values range from 0 to 1, where 1 indicates perfectly homogeneous labelling. Notice that by switching true cluster labels with the predicted cluster labels, the Completeness Score is obtained.

#### Completeness Score

The result of the tested clustering approach satisfies the Completeness Score (CS) [52] if all the cells that belong to a given cell-type are elements of the same cluster. Its values range from 0 to 1, where 1 indicates perfectly complete labelling. Notice that by switching true cluster labels with the predicted cluster labels, the HS is obtained.

#### V-Measure

The V-Measure (VM) [53] is the harmonic mean between HS and CS; it is equivalent to MII when the arithmetic mean is used as aggregation function.

### Integration of multiple datasets obtained with the same sequencing platforms

Nowadays, various scRNA-Seq platforms are currently available (e.g., droplet-based and plate-based [54–63]) and their integration is often challenging due to the differences in biological sample batches as well as to the used experimental platforms. To test whether AEs can be effectively applied to combine multiple datasets, generated using the same platform but under different experimental conditions, we used the PBMC datasets.

We merged the control and treated datasets by first using vanilla PCA and AEs, and then PCA and AEs followed by either BBKNN or Harmony, ComBat, and CCA. After the construction of the neighbourhood graphs, we performed a clustering step by using the Leiden algorithm [64]. Since in the original paper 8 different cell-types were manually identified [41], we selected Leiden’s resolutions that allowed us to obtain 8 distinct clusters and calculated all the metrics described above. In what follows, the calculated values of all metrics are given in percentages (mean ± standard deviation). For each metric, the higher the value the better the result.

Our analysis showed that the CCA-based approach, proposed in the Seurat library, achieved a mean ARI equal to 73.49% (±1.52%), ComBat reached a mean ARI of 72.84% (±0.78%), vanilla PCA had a mean ARI of 68.90% (±0.86%), PCA followed by BBKNN was able to obtain a mean ARI of 83.65% (±0.81%), while PCA followed by Harmony reached a mean ARI of 82.83% (±1.20%), as shown in Figure 3A. Among all the tested AEs, MMDAE followed by Harmony (using the NB loss function, 256 neurons for the hidden layer, and 32 neurons for the latent space) achieved the best results, with a mean ARI equal to 87.18% (±0.49%). In order to assess whether any of the results obtained by the best AE were different from a statistical point of view, we applied the Mann–Whitney U test with the Bonferroni correction [65–67]. In all the comparisons, MMDAE followed by Harmony had a p-value lower than 0.0001, confirming that the achieved results are statistically different compared to those achieved by the other approaches.

**Figure 2.**
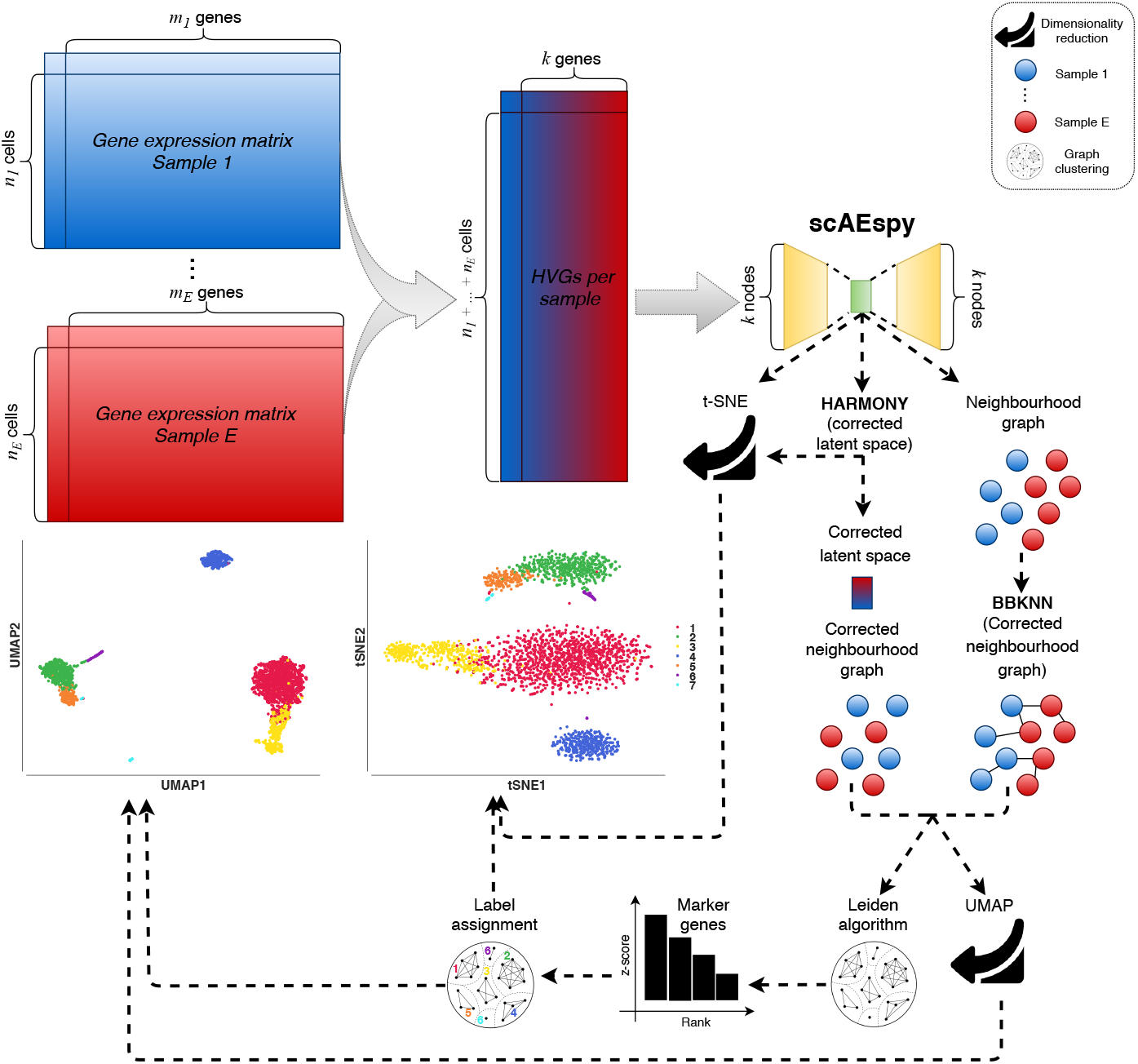
The proposed workflow to integrate different samples. Given *E* different samples, their gene expression matrices are merged. Then, the top *k* HVGs are selected by considering the different samples. Specifically, they are selected within each sample separately and then merged to avoid the selection of batch-specific genes. scAEspy is used to reduce the HVG space (*k* dimensions), and the obtained latent space can be (*i*) used to calculate a t-SNE space, (*ii*) corrected by Harmony, and (*iii*) used to infer an uncorrected neighbourhood graph. The corrected latent space by Harmony is then used to build a neighbourhood graph, which is clustered by using the Leiden algorithm and used to calculate a UMAP space. Otherwise, BBKNN is applied to rebuild a uncorrected neighbourhood graph by taking into account the possible batch-effects. The corrected neighbourhood graph by BBKNN is then clustered by using the Leiden algorithm and used to calculate a UMAP space. In order to assign the correct label to the obtained clusters, the marker genes are calculated by using the Mann–Whitney U test. Finally, the annotated clusters can be visualised in both t-SNE and UMAP space.

**Figure 3.**
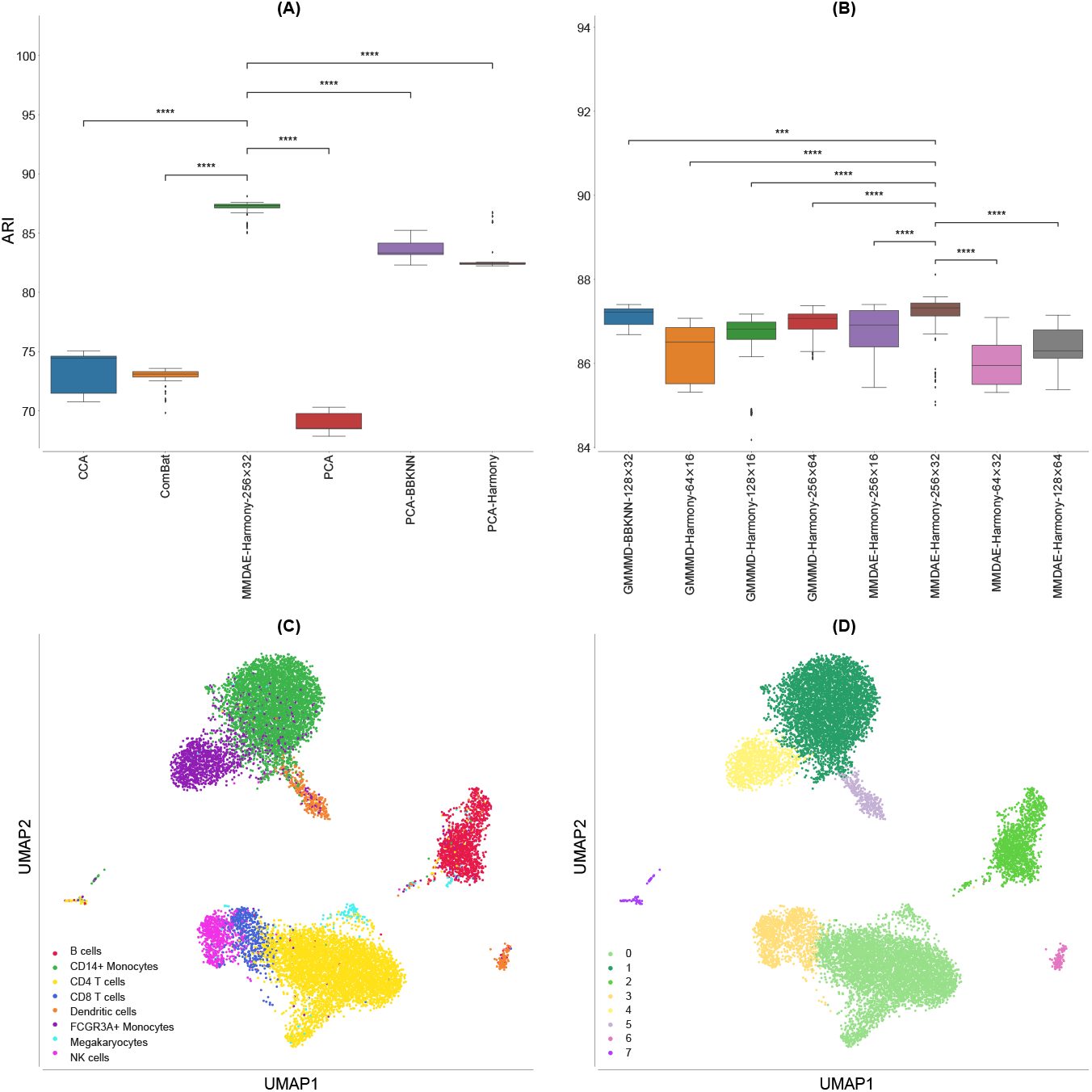
Results obtained on the PBMC datasets. **(A)** Boxplot showing the ARI values achieved by CCA, ComBat, PCA, MMDAE followed by Harmony with dimension (256, 32), PCA followed by BBKNN, and PCA followed by Harmony on the PBMC datasets. **(B)** Boxplot showing the ARI values achieved by the best AE for each of the tested dimension (*H, L*) of the hidden layer (*H* neurons) and latent space (*L* neurons). **(C)** UMAP visualisation of the cell-type manually annotated in the original paper. **(D)** UMAP visualisation of clusters identified by the Leiden algorithm using the resolution corresponding by the best ARI achieved by MMDAE followed by Harmony. p-value≤ 0.0001 (****); 0.0001 <p-value≤ 0.001 (***); 0.001 <p-value≤ 0.01 (**); 0.01 <p-value≤ 0.05 (*); p-value*>* 0.05 (ns)

Regarding the AMII, CCA had a mean value of 66.46% (±0.50%), ComBat achieved a mean value of 70.95% (±0.82%), vanilla PCA obtained a mean value of 68.44% (±1.00%), PCA followed by BBKNN reached a mean value of 75.22% (±0.76%), while PCA followed by Harmony reached a mean value of 74.55% (±1.19%). MMDAE followed by Harmony had better results, with a mean value equal to 78.61% (±0.29%). MMDAE followed by Harmony outperformed the other strategies also in terms of of FMS, HS, CS, and VM (see Additional file 2 and Figure 4).

**Figure 4.**
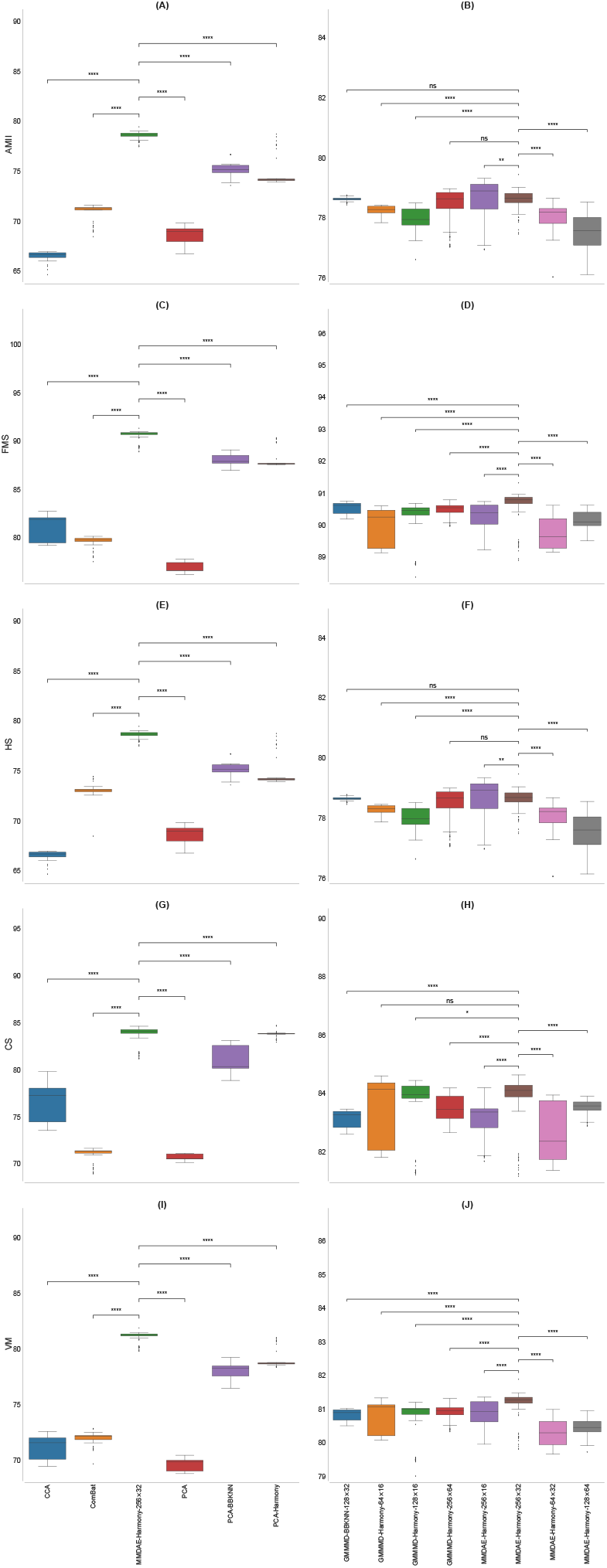
Boxplot showing the values of the calculated metrics using CCA, ComBat, PCA, MMDAE followed by Harmony with dimension (256, 32), PCA followed by BBKNN, and PCA followed by Harmony as well as by the best AE for each of the tested dimension (*H, L*), analysing the PBMC datasets. **(A)** AMII achieved by the different strategies. **(B)** AMII achieved by the best AE for each of the tested dimension. **(C)** FMS achieved by the different strategies. **(D)** FMS achieved by the best AE for each of the tested dimension. **(E)** HS achieved by the different strategies. **(F)** HS achieved by the best AE for each of the tested dimension. **(G)** CS achieved by the different strategies. **(H)** CS achieved by the best AE for each of the tested dimension. **(I)** VM achieved by the different strategies. **(J)** VM achieved by the best AE for each of the tested dimension. p-value≤ 0.0001 (****); 0.0001 <p-value≤ 0.001 (***); 0.001 <p-value≤ 0.01 (**); 0.01 <p-value≤ 0.05 (*); p-value*>* 0.05 (ns)

We also compared the results obtained by the best AE for each of the tested dimension (*H, L*) in terms of ARI (Figure 3B). GMMMD followed by Harmony (using the NB loss function) obtained the best results for the dimension (64, 16), GMMMD followed by Harmony (using the Poisson loss function) reached the best results for the dimensions (128, 16) and (256, 64), and GMMMD followed by BBKNN (using the NB loss function) achieved the best results for the dimension (128, 32). MMDAE followed by Harmony (using the NB loss function) was able to reach the best results for the dimensions (64, 32), (256, 16), and (256, 32), while MMDAE followed by Harmony (using the Poisson loss function) obtained the best result for the dimensions (128, 64). Notice that we used two Gaussian distributions because we merged two different datasets.

In order to visually assess the quality of the separation of the manually annotated cell-type and the found clusters, we plotted them in the UMAP space generated starting from the MMDAE followed by Harmony space (Figures 3C and D). Finally, we also plotted the two samples in the same UMAP space to visually see the quality of the alignment between the two samples themselves (Figure 5A). This plot confirms that the batch-effects were completely removed.

**Figure 5.**
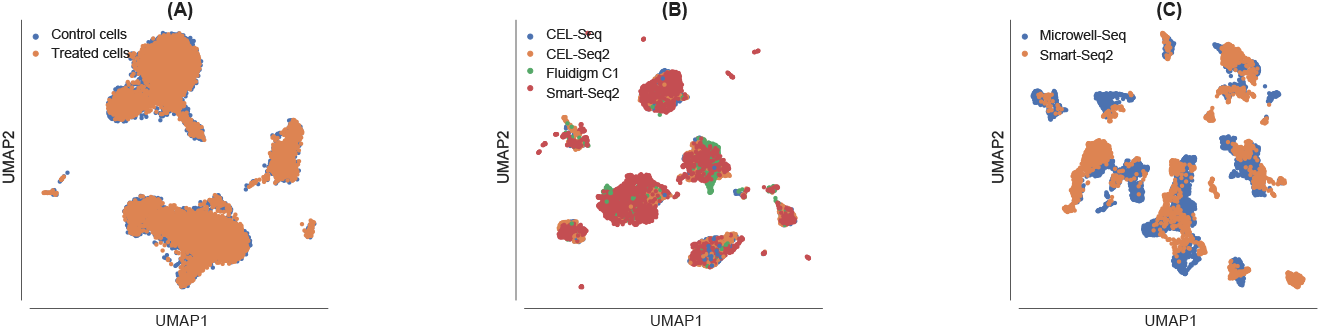
UMAP visualisation showing the sample alignment performed by Harmony into the latent space obtained by MMDAE with dimension (256, 32) for the PBMC datasets **(A)**, by GMMMD with dimension (256, 32) for the PIC datasets **(B)**, and by MMDVAE with dimension (256, 64) for the MCA datasets **(C)**.

Our analysis showed that clustering the neighbourhood graph generated from AE spaces allowed for a better identification of the existing cell-types when compared to other approaches, thus confirming the ARI results.

### Integration of multiple datasets obtained with different sequencing platforms

Combining datasets from different studies and scRNA-Seq platforms can be a powerful approach to obtain complete information about the biological system under investigation. However, when datasets generated with different platforms are combined, the high variability in the gene expression matrices can obscure the existing biological relationships. For example, the gene expression values are much higher in data acquired with plate-based methods (i.e., up to millions) than in those acquired with droplet-based methods (i.e., a few thousands). Thus, combining gene expression data that spread across several orders of magnitude is a difficult task that cannot be tackled by using linear approaches like PCA. To examine how well AEs perform in solving this issue, we combined four PIC datasets acquired with CEL-Seq [56], CEL-Seq2 [57], Fluidigm C1 [63], and Smart-Seq2 protocols [62].

We integrated the datasets by first using vanilla PCA and AEs, and then PCA and AEs followed by either BBKNN or Harmony, ComBat, and CCA. Since in the original paper 13 cell-types were manually annotated for the PIC datasets [46], we clustered the neighbourhood graphs using the Leiden algorithm considering only the resolutions that allowed us to obtain 13 distinct clusters. We then calculated ARI, AMII, FMS, HS, CS, and VM metrics. The calculated values of all metrics are given in percentages (mean ± standard deviation); for each metric, the higher the value the better the result.

CCA had a very low mean ARI, i.e., 5.45% (±0.22%), ComBat obtained a mean ARI of 76.20% (±3.06%), vanilla PCA achieved a mean ARI of 61.38% (±0.09%), PCA followed by BBKNN reached a mean ARI of 71.49% (±0.58%), while PCA followed by Harmony was able to obtain a mean ARI of 94.00% (±0.36%), see Figure 6A. GMMMD followed by Harmony (using the NB loss function, 256 neurons for the hidden layer, and 32 neurons for the latent space) outperformed the other AEs, achieving a mean ARI equal to 94.23% (±0.12%). In all the comparisons, except for the one against PCA followed by Harmony, GMMMD followed by Harmony had a p-value lower than 0.0001, confirming that the achieved results are statistically different with respect to those obtained by the other approaches.

**Figure 6.**
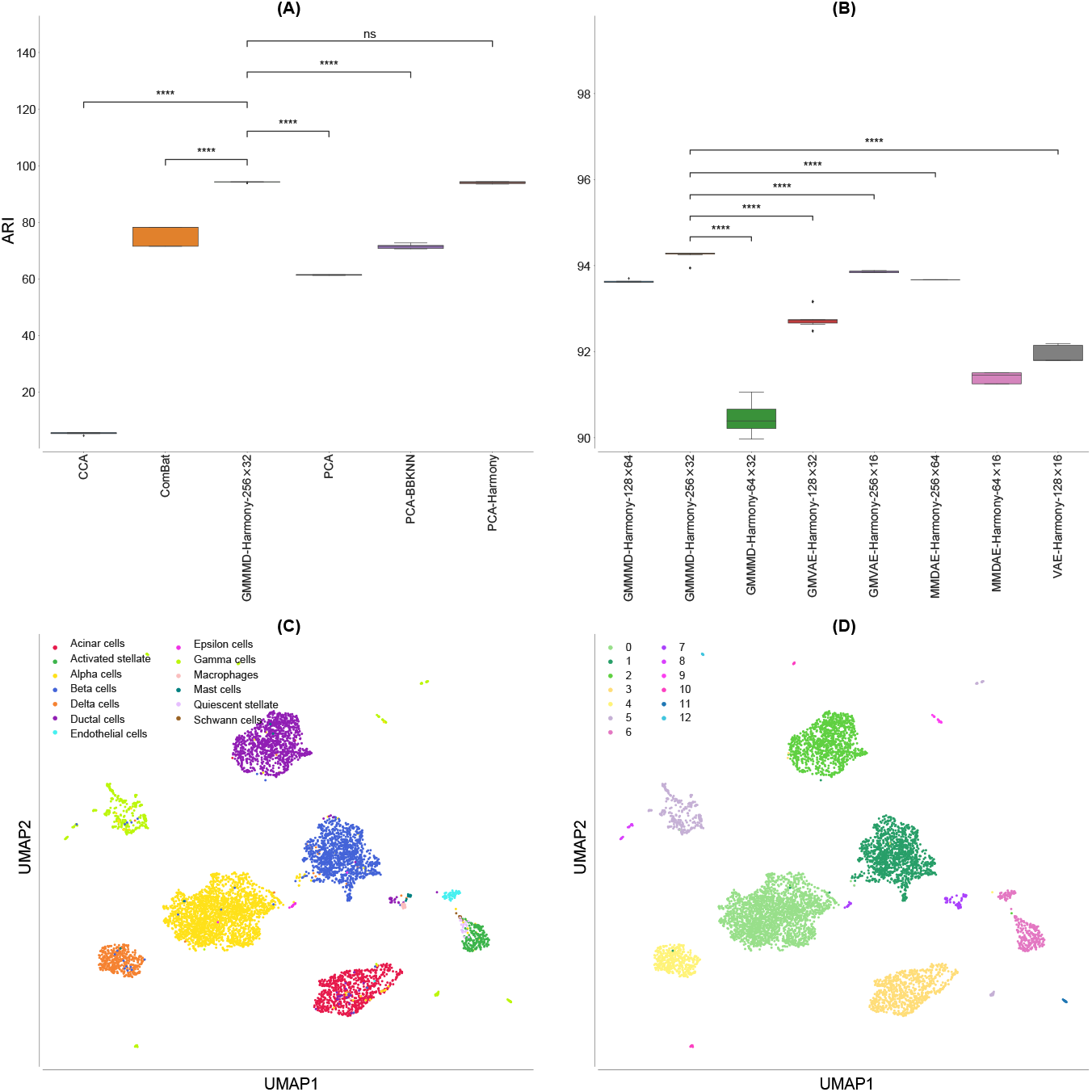
Results obtained on the PIC datasets. **(A)** Boxplot showing the ARI values achieved by CCA, ComBat, PCA, GMMMD followed by Harmony with dimension (256, 32), PCA followed by BBKNN, and PCA followed by Harmony on the PBMC datasets. **(B)** Boxplot showing the ARI values achieved by the best AE for each of the tested dimension (*H, L*) of the hidden layer (*H* neurons) and latent space (*L* neurons). **(C)** UMAP visualisation of the cell-type manually annotated in the original paper. **(D)** UMAP visualisation of clusters identified by the Leiden algorithm using the resolution corresponding by the best ARI achieved by GMMMD followed by Harmony. p-value≤ 0.0001 (****); 0.0001 <p-value≤ 0.001 (***); 0.001 <p-value≤ 0.01 (**); 0.01 <p-value≤ 0.05 (*); p-value*>* 0.05 (ns)

Similar results were achieved for the AMII metric: CCA reached a mean value equal to 16.57% (±0.33%), ComBat obtained a mean value of 76.11% (±1.14%), vanilla PCA reached a mean value of 71.70% (±0.11%), PCA followed by BBKNN achieved a mean value of 77.55% (±0.65%) and PCA followed by Harmony a mean value of 91.17% (±0.47%), while GMMMD followed by Harmony was able to reach a mean value equal to 89.37% (±0.02%). Considering the other measures, both PCA and GMMMD followed by Harmony obtained very similar results, outperforming the other strategies (see Additional file 3 and Figure 7).

**Figure 7.**
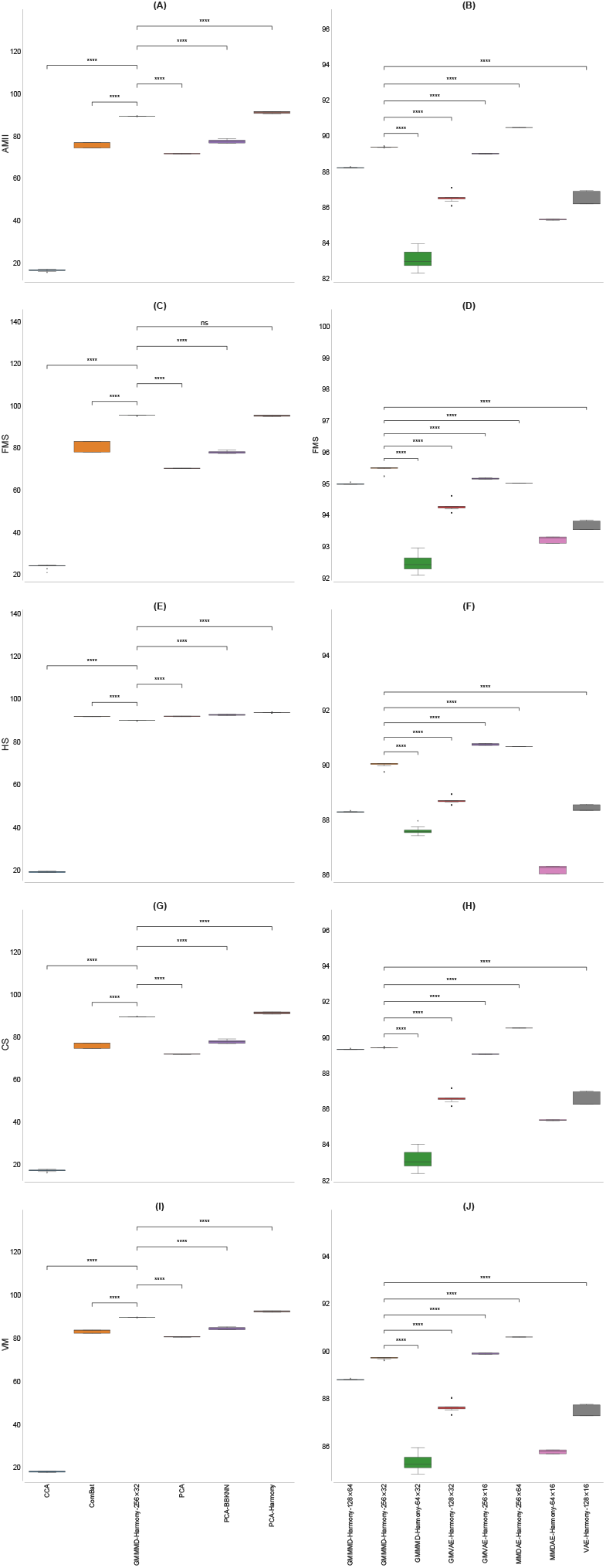
Boxplot showing the values of the calculated metrics using CCA, ComBat, PCA, GMMMD followed by Harmony with dimension (256, 32), PCA followed by BBKNN, and PCA followed by Harmony as well as by the best AE for each of the tested dimension (*H, L*), analysing the PIC datasets. **(A)** AMII achieved by the different strategies. **(B)** AMII achieved by the best AE for each of the tested dimension. **(C)** FMS achieved by the different strategies. **(D)** FMS achieved by the best AE for each of the tested dimension. **(E)** HS achieved by the different strategies. **(F)** HS achieved by the best AE for each of the tested dimension. **(G)** CS achieved by the different strategies. **(H)** CS achieved by the best AE for each of the tested dimension. **(I)** VM achieved by the different strategies. **(J)** VM achieved by the best AE for each of the tested dimension. p-value≤ 0.0001 (****); 0.0001 <p-value≤ 0.001 (***); 0.001 <p-value≤ 0.01 (**); 0.01 <p-value≤ 0.05 (*); p-value*>* 0.05 (ns)

Considering the best AE for each of the tested dimension (*H, L*) in terms of ARI (see Figure 6B), GMMMD followed by Harmony (using the NB loss function) resulted the best choice for the dimensions (128, 64) and (256, 32), while it obtained the best results for the dimension (64, 32) when the Poisson loss function was used. GMVAE followed by Harmony (using the Poisson loss function) reached the best results for the dimensions (128, 32) and (256, 16). MMDAE followed by Harmony achieved the best results for the dimensions (256, 64) and (64, 16), exploiting the NB loss function and Poisson loss function, respectively. Finally, VAE followed by Harmony obtained the best results with the Poisson function for the dimension (128, 16). Note that we exploited four Gaussian distributions because we merged four different datasets.

The quality of the separation of the manually annotated cell-type and found clusters can be visually evaluated in Figures 6C and D. We finally visualised the cells (coloured by platform) using the UMAP space generated from the GMMMD followed by Harmony space (Figure 5B) to confirm that the batch-effects among the samples sequenced with different platforms were correctly removed. Taken together, our analysis shows that GMMMD followed by Harmony can efficiently identify the “shared” cell-types across the different platforms due to its ability to deal with the high variability in the gene expression matrices. We would like to highlight that PCA followed by Harmony was capable of achieving good results because the original clusters were obtained by applying a similar pipeline [46].

As a final test, we combined two MCA datasets acquired with Microwell-Seq [47] and Smart-Seq2 protocols [62]. We integrated the datasets in the same way we did in the other two tests. We clustered the neighbourhood graphs using the Leiden algorithm considering only the resolutions that allowed us to obtain 11 distinct clusters because 11 distinct cell-types were manually annotated for the PIC datasets [47]. We then calculated all metrics.

In such a case, MMDVAE followed by Harmony (using the Poisson loss function, 256 neurons for the hidden layer, and 64 neurons for the latent space) outperformed the other AEs as well as the other strategies, obtaining a mean ARI equal to 79.50% (±0.02%), as shown in Figure 8A. ComBat achieved the worst mean ARI, i.e., 54.13% (±4.22%), CCA reached a mean ARI of 57.62% (±0.75%), vanilla PCA obtained a mean ARI of 67.29% (±11.55%), PCA followed by BBKNN had a similar mean ARI, that is, 67.73% (±3.98%), while PCA followed by Harmony achieved a mean ARI of 66.08% (±0.11%). MMDVAE followed by Harmony had a p-value lower than 0.0001 in all the tested comparisons. Considering the other metrics, MMDVAE followed by Harmony generally obtained better results compared to the other strategies (see Additional file 3, Figure 8B, and Figure 9).

**Figure 8.**
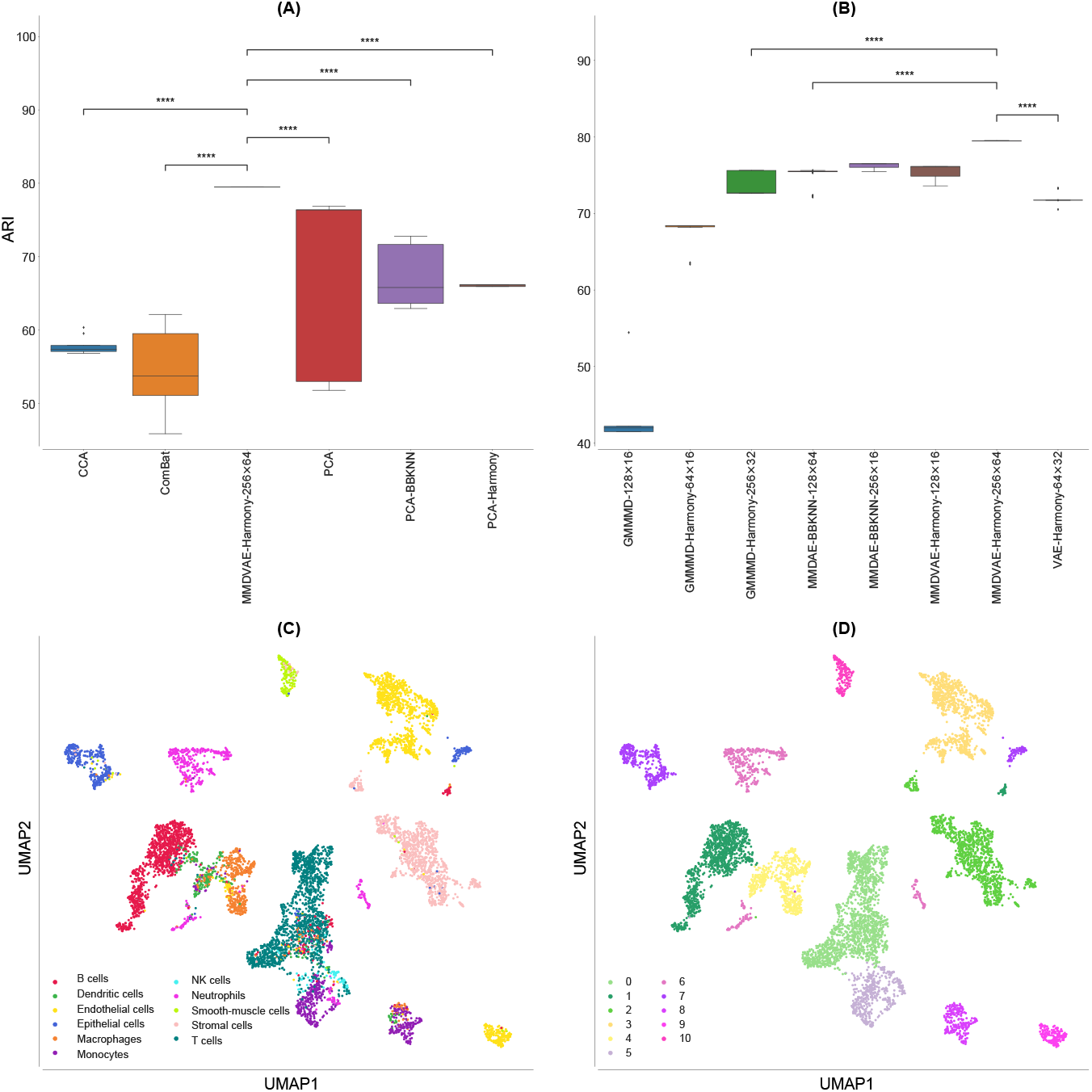
Results obtained on the MCA datasets. **(A)** Boxplot showing the ARI values achieved by CCA, ComBat, PCA, MMDVAE followed by Harmony with dimension (256, 64), PCA followed by BBKNN, and PCA followed by Harmony. **(B)** Boxplot showing the ARI values achieved by the best AE for each of the tested dimension (*H, L*) of the hidden layer (*H* neurons) and latent space (*L* neurons). **(C)** UMAP visualisation of the cell-type manually annotated in the original paper. UMAP visualisation of clusters identified by the Leiden algorithm using the resolution corresponding by the best ARI achieved by MMDVAE followed by Harmony. p-value 0.0001 (****); 0.0001 <p-value 0.001 (***); 0.001 <p-value ≤ 0.01 (**); 0.01 <p-value ≤ 0.05 (*); p-value*>* ≤ 0.05 (ns)

**Figure 9.**
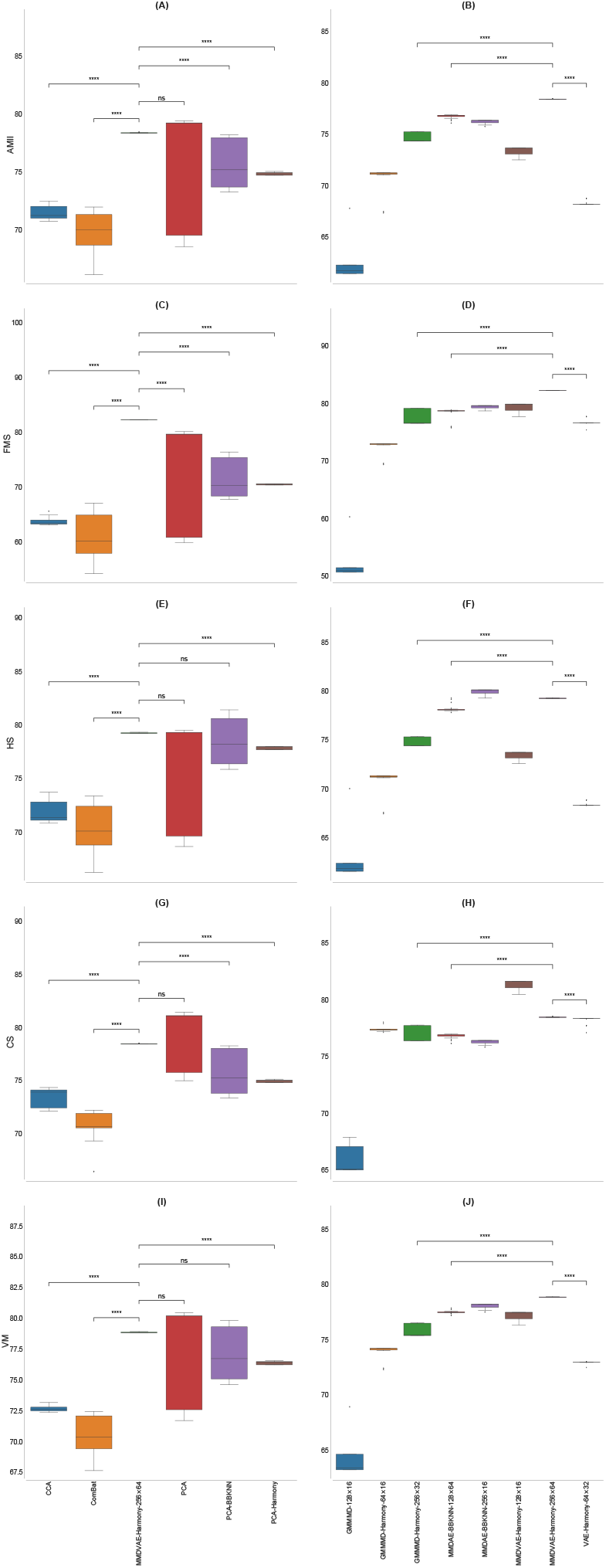
Boxplot showing the values of the calculated metrics using CCA, ComBat, PCA, MMDVAE followed by Harmony with dimension (256, 64), PCA followed by BBKNN, and PCA followed by Harmony, as well as by the best AE for each of the tested dimension (*H, L*), analysing the MCA datasets. **(A)** AMII achieved by the different strategies. **(B)** AMII achieved by the best AE for each of the tested dimension. **(C)** FMS achieved by the different strategies. **(D)** FMS achieved by the best AE for each of the tested dimension. **(E)** HS achieved by the different strategies. **(F)** HS achieved by the best AE for each of the tested dimension. **(G)** CS achieved by the different strategies. **(H)** CS achieved by the best AE for each of the tested dimension. **(I)** VM achieved by the different strategies. **(J)** VM achieved by the best AE for each of the tested dimension. p-value≤ 0.0001 (****); 0.0001 <p-value≤ 0.001 (***); 0.001 <p-value≤ 0.01 (**); 0.01 <p-value≤ 0.05 (*); p-value*>* 0.05 (ns)

Comparing the best AE for each of the tested dimension (*H, L*) in terms of ARI, the vanilla GMMMD with the NB loss function obtained the best results for the dimension (128, 16), while GMMMD followed by Harmony reached the best results for the dimensions (256, 32) and (64, 16), exploiting the Poisson loss function and NB loss function, respectively. MMDAE followed by BBKNN (using the ZIP loss function) achieved the best results for the dimensions (256, 16) and (128, 64), exploiting the NB loss function and Poisson loss function, respectively. MMDVAE followed by Harmony resulted the best choice for the dimensions (128, 16) and (256, 64) when coupled with the NB loss function and Poisson loss function, respectively. Finally, VAE followed by Harmony with the Poisson loss function obtained the best results for the dimension (64, 32). As for the the integration of the PBMC datasets, we used two Gaussian distributions because we merged two different datasets.

Figures 8C and D show the UMAP generated from the MMDVAE followed by Harmony space coloured by the manually annotated cell-type and found clusters, respectively, while Figure 5C depicts the cells coloured by platform on the same UMAP space, confirming that the batch-effects between the two samples were correctly removed. In this case, the achieved results show that MMDVAE followed by Harmony was able to better identify the “shared” cell-types across the different platforms.

In order to test the computational performance of scAEspy, we considered a VAE with a single hidden layer composed of 256 neurons, a latent space of 32 neurons, and the NB loss function. All tests were executed on a laptop equipped with an Intel Core i7-1165G7 CPU (clock 2.8 GHz) and 16 GB of RAM, running on Ubuntu 20.04 LTS. In all the tested datasets, we used the top 1000 HVGs. The executions lasted 95.02 seconds, 101.05 seconds, and 223.45 seconds for PIC, MCA, and PBMC, respectively. Considering that PIC, MCA, and PBMC are composed of 6321, 6954, and 14039 cells, respectively, scAEspy scales linearly with the number of cells.

## Discussion

Non-linear approaches for dimensionality reduction can be effectively used to capture the non-linearities among gene interactions that may exist in the high-dimensional expression space of scRNA-Seq data [15]. Among the different non-linear approaches, AEs showed outstanding performance, outperforming other approaches like UMAP and t-SNE. Several AE-based methods have been developed so far, but their integration with the common single-cell toolkits results in a difficult task because they usually require input data codified in a specific format. In addition, three different machine learning libraries are required to run them (i.e., Keras, TensorFlow, PyTorch).

Here, we proposed scAEspy, a unifying and user-friendly tool that allows the user to use the most recent and promising AEs (i.e., VAE, MMDAE, MMDVAE, and GMVAE). We also designed and developed GMMMD and GMMMDVAE, two novel AEs that combine MMDAE and MMDVAE with GMVAE to exploit more than one Gaussian distribution. We introduced a learnable prior distribution in the latent space to model the dimensionality of the subpopulations of cells composing the data, or to combine multiple samples.

We integrated AEs with both Harmony and BBKNN to remove the existing batch-effects among different datasets. Our results showed that exploiting the latent space to remove the existing batch-effects permits for better identification of the cell sub-populations. As a batch-effect removal tool, Harmony allowed for achieving better results than BBKNN in the majority of the cases. When different droplet-based data have to be combined, our GMMMD and the MMDAE, coupled with the constrained NB and Poisson loss functions, obtained the best results compared to all the other AEs. In order to combine and analyse multiple datasets, generated by using different scRNA-Seq platforms, both GMMMD and MMDVAE, mainly used together with the NB and Poisson loss functions, outperformed the other strategies. However, also GMVAE and the simple VAE obtained outstanding performance, highlighting that the Kullback-Leibler divergence function can become fundamental to handle data spreading various orders of magnitude, especially the high values (up to millions) introduced by plate-based methods. It is clear that using more than one Gaussian distribution allows for obtaining a better integration of the datasets and separation of the cell-types when more than two datasets have to be integrated, as clearly shown by the results reached on the PIC datasets.

As a good practice, we suggest filtering the gene expression matrices using the top HVGs (i.e., 1000), calculated separately within each batch and merged to avoid the selection of batch-specific genes. The original counts, corresponding to the top HVGs, should be the input data of scAEspy. GMMMD coupled with Harmony should be used to integrate different droplet-based datasets as well as to merge datasets generated by using different scRNA-Seq platforms. Harmony should be applied to the latent space produced by GMMMD to take into account the possible batches among the datasets. When a single dataset has to be analysed, the users should use MMDAE with droplet-based data, while MMDVAE should be preferred for data acquired with plate-based methods. In all cases, using a single hidden layer composed of 256 neurons, a latent space with 32 neurons, and the constrained NB or Poisson loss function generally allows for obtaining satisfactory results.

Considering the achieved results on the identification of the clusters, scAEspy can be used at the basis of methods that aim at automatically identifying the cell-types composing the scRNA-Seq datasets under analysis [68]. As a matter of fact, scAEspy coupled with BBKNN was successfully applied to integrate 15 different foetal human samples, enabling the identification of rare blood progenitor cells [69].

## Conclusions

In this study, we proposed an AE-based and user-friendly tool, named scAEspy, which allows for using the most recent and promising AEs to analyse scRNA-Seq data. The user can select the desired AE by only setting up two user-defined parameters. Once the selected AE has been trained, it can be used to generate synthetic cells to increase the number of data for further downstream analyses (e.g., training classifiers). In scAEspy, the latent space is easily accessible and thus allows the user to perform different types of analysis, such as the correction of possible batch-effect in a reduced non-linear space or the inference of differentiation trajectories. In this case, the latent space can be utilised to generate the “pseudotime” that measures transcriptional changes that a cell undergoes during the dynamic process.

Thanks to its modularity, scAEspy can be extended to accommodate new AEs so that the user will be always able to utilise the latest and cutting-edge AEs [70], which can improve the downstream analysis of scRNA-Seq data. It is worth noticing that scAEspy can be used on HPC infrastructures, both based on CPUs and GPUs, to speed-up the computations. This is a crucial point when datasets composed of hundreds of thousands of cells are analysed. In such cases, the required running time drastically increases, so relying on HPC infrastructures is the best solution to incredibly reduce the prohibitive running time.

### Future improvements

As an improvement, prior biological knowledge about genes from ontologies can be incorporated into scAEspy. Ontologies can introduce useful information into machine learning systems that are used to solve biological problems. They allow for integrating data from different omics (e.g., genomics, transcriptomics, proteomics, and metabolomics) as structured representations of semantic knowledge, which is commonly used for the representation of biological concepts. This approach has been successfully applied to predict the clinical targets from high-dimensional low-sample data [71]. Specifically, ontology embeddings are able to capture the semantic similarities among the genes, which can be exploited to sparsify the network connections. In addition, the Gene Ontology (GO) [72] can be exploited to interpret the extracted features from the latent spaces generated by the AEs, thus bringing an explanation to the learned representations of the gene expression data. As a possible example, g:Profiler [73] focusing on GO terms, Kyoto Encyclopedia of Genes and Genomes (KEGG), and Reactome can be used on the learned embeddings to investigate the joint effects of different gene sets within specific biological pathways. This approach can help the interpretability and explainability of the learned embeddings of the used AEs.

### Integration of multi-omics data

Since AEs showed outstanding performance in the integration of multi-omics of cancer data [70], we plan to extend scAEspy to analyse other single-cell omics. For instance, AEs can be applied to analyse scATAC-seq, where the identification of the cell-types is still very difficult due to technical challenges [10, 74]. scAEspy could be effectively applied to analyse disparate types of single-cell data from different points of view. The latent representations of different or combined single-cell omics can be used for further and more in-depth analyses. For instance, the application of other machine learning techniques (e.g., deep neural networks) to the latent representations could facilitate the identification of interesting patterns on gene expression or methylation data, as well as relationships among genomics variants. In that regard, scAEspy can be the starting point to build a more comprehensive toolkit designed to integrate multi single-cell omics as an integration and extension of the work proposed in [70].

## Methods

We developed scAEspy so that it can be easily integrated into both Scanpy and Seurat pipelines, as it directly works on a gene expression matrix (see Figure 2). We integrated into a single tool the latest and most powerful AEs designed to resolve the problems underlying scRNA-Seq data (e.g., sparsity, intrinsic noise, dropout events [3]). Specifically, scAEspy is comprised of six AEs, based on the VAE [17] and InfoVAE [35] architectures. The following most advanced AEs are included in scAEspy: VAE, MMDAE, MMDVAE, GMVAE, and two novel Gaussian-mixture AEs that we developed, called GMMMD and GMMMDVAE. GMMMD is a modification of the MMDAE where more than one Gaussian distribution is used to model different modes and only the MMD function is used as divergence function. GMMMDVAE is a combination of MMDVAE and GMVAE where both the MMD function [75] and the Kullback-Leibler divergence function [76] are used. scAEspy allows the user to exploit these six different AEs by setting up two user-defined parameters, *α* and *λ*, which are needed to balance the MMD and the Kullback-Leibler divergence functions. We designed and developed GMMMD and GMMMDVAE starting from InfoVAE [35] and scVAE [20]. In addition, a learnable mixture distribution was used for the prior distribution in the latent space, and also the marginal conditional distribution was defined to be a learnable mixture distribution with the same number of components as the prior distribution. Finally, the user can also select the following loss functions: NB, constrained NB, Poisson, constrained Poisson, ZINB, constrained ZINB, ZIP, constrained ZIP, and Mean Square Error (MSE).

### The tested batch-effect removal tools

Originally proposed to deal with batch-effects in microarray gene expression data [38], ComBat has been successfully applied to analyse scRNA-Seq data [77]. Briefly, given a gene expression matrix, it is firstly standardised so that all genes have similar means and variances. Then, starting from the obtained standardised matrix, standard distributions are fitted using a Bayesian approach to estimate the existing batch-effects in the data. Finally, the original expression matrix is corrected using the computed batch-effect estimators. In our tests, we used the default parameter settings provided by the Scanpy function combat. We then applied PCA on the space obtained by the top *k* (here, we set *k* = 1000) HVGs calculated by using the function provided by Scanpy (v.1.4.5.1), where the top HVGs are separately selected within each batch and merged to avoid the selection of batch-specific genes. We calculated the first 50 components and applied the so-called “elbow method” to select the number of components for the downstream analysis [8]. The “elbow method” was applied taking into consideration the variance explained by each PCA component, which can be visualised using the Scanpy function pca_variance_ratio. Including less informative PCA components might help in the identification of rare cell types, but they can also introduce noise that hinders the downstream analysis steps; thus, we opted to include only the most informative PCA components. Specifically, we used the first 18, 13, and 18 components for PBMC, PIC, and MCA datasets, respectively. After that, we calculated the neighbourhood graph by using the default parameter settings proposed in Scanpy. We clustered the obtained neighbourhood graphs with the Leiden algorithm by selecting the values of the resolution parameter such that the number of clusters was equal to the manually annotated clusters. Finally, all the metrics for each found resolution have been calculated.

As another batch-effect removal tool, we used the CCA-based approach proposed in the Seurat package (v.2.3.4) [13]. We applied both RunCCA and MultiCCA Seurat functions to integrate two batches and more than two batches, respectively. Firstly, we normalised and log-transformed the counts. Then, we calculated the top 1000 HVGs by using the function provided by Scanpy (v.1.4.5.1). We also scaled the log-transformed data to zero mean and unit variance. In both RunCCA and MultiCCA Seurat functions, as a first step, the CCA components were exploited to compute the linear combinations of the genes with the maximum correlation between the batches. We used the default number of CCA components (i.e., 20) provided by the functions RunCCA and MultiCCA of the Seurat package, as also suggested in [36]. A dynamic time warping (AlignSubspace Seurat function), which accounts for population density changes, was then used to align the calculated vectors and obtain a single low-dimensional subspace where the batch-effects are corrected. We calculated the neighbourhood graph, using the default parameter settings proposed in Scanpy, starting from the aligned low-dimensional subspace. We clustered the built neighbourhood graphs with the Leiden algorithm as explained before. Finally, we calculated all the metrics for each found resolution.

We also applied Harmony [37] to remove the batch-effects. Starting from a reduced space (e.g., PCA space or latent space), Harmony exploits an iterative clustering-based procedure to remove the multiple-dataset-specific batch-effects. In each iteration, the following four steps are applied: (*i*) the cells are grouped into multiple-dataset clusters by exploiting a variant of the soft *k*-means clustering, which is a fast and flexible method developed to cluster single-cell data; (*ii*) a centroid is calculated for each cluster and for each specific dataset; (*iii*) using the calculated centroids, a correction factor is derived for each dataset; (*iv*) the correction factors are then used to correct each cell with a cell-specific factor.

As a further batch-effect removal tool, we applied BBKNN [36]. Polanski *et al*. [36] showed that BBKNN has comparable or better performance in removing batch-effects with respect to the CCA-based approach proposed in the Seurat package, Scanorama [78], and mnnCorrect [79]. In addition, BBKNN is a lightweight graph alignment method that requires minimal changes to the classical workflow. Indeed, it computes the *k*-nearest neighbours in a reduced space (e.g., PCA or latent space), where the nearest neighbours are identified in a batch-balanced manner using a user-defined distance (in our tests, we used the Euclidean distance). The neighbour information is transformed into connectivities to build a graph where all cells across batches are linked together. We used both Harmony and BBKNN to correct the PCA and AE spaces.

As a final step, we calculated the UMAP spaces starting from the built neighbourhood graphs and using the default parameter settings proposed in Scanpy, except for the initialisation of the low dimensional embedding (i.e., *init_pos* equal to *random*, and *random_state* equal to 10 of the umap function).

### The proposed pipeline

We modified the workflow shown in Figure 1 by replacing PCA with AEs (Figure 2). We merged the gene expression matrices of *E* different samples (*E* = 2, *E* = 4, and *E* = 2 for PBMC, PIC, and MCA datasets, respectively). We applied both PCA and AEs on the space obtained by the top 1000 HVGs calculated by using the implementation provided by Scanpy (v.1.4.5.1).

For what concerns PCA, we firstly normalised and log-transformed the counts, then we applied a classic standardisation, that is, the distribution of the expression of each gene was scaled to zero mean and unit variance. We calculated the first 50 components; after that, we used the “elbow method” to select the first 12, 14, and 19 components for the PBMC, PIC, and MCA datasets, respectively. As in the case of the ComBat workflow, the “elbow method” was applied considering the variance explained by each PCA component.

Regarding AEs, we used the original counts since AEs showed to achieve better results when applied using the raw counts [20]. Indeed, using the counts allows for exploiting discrete probability distributions, such as Poisson and NB distributions, which obtained the best results in our tests. In all the tests presented here, we used a single hidden layer. In addition, we set 100 epochs, sigmoid activation functions, and a batch equal to 100 samples (i.e., cells). In all tests we used the Adam optimizer [80]. After that, we applied three different strategies (Figure 2): (*i*) we calculated the neighbourhood graph in both PCA and AE spaces by using the default parameter settings proposed in Scanpy. Then, we clustered the obtained neighbourhood graphs with the Leiden algorithm as described before. Finally, we calculated all the metrics for each found resolution; (*ii*) we performed a similar analysis where we firstly corrected the PCA and AE spaces using Harmony [37] with the default parameter settings proposed in https://github.com/slowkow/harmonypy; (*iii*) we performed the same analysis described in (*i*) by replacing the neighbourhood graphs with those generated using BBKNN, using the default parameter settings.

It is worth mentioning that in all the tested batch-effect removal tools, we used different samples that have to be merged as batches. Specifically, the number of batches is equal to 2, 4, and 2 for the PBMC, PIC, and MCA datasets, respectively.

### The generalised formulation of scAEspy

In this work, we used the notation proposed in [35] to extend MMDVAE with multiple Gaussian distributions as well as to introduce a learnable prior distribution in the latent space. The idea behind the introduction of learnable coefficients is that they might be suitable to model the diversity among the subpopulations of cells composing the data or to combine multiple samples or datasets.

We consider *p*^*\**^(**x**) as the unknown probability in the input space over which the optimisation problem is formulated, ***z*** is the latent representation of ***x*** with

|***z***| ≤ |***x***|. The encoder is identified by a function *e*_***φ***_ : ***x*** *→* ***z***, while the decoder by a function *d*_***θ***_ : ***z*** *→* ***x***.

We remind that in VAEs, the input ***x*** is not mapped into a single point in the latent space, but it is represented by a probability distribution over the latent space. In what follows, we denote by *q*(***z***) any possible distribution in the latent space and by *y ∈ {*1, …, *K}* a categorical random variable, where *K* corresponds to the number of desired Gaussian distributions. As general strict divergence function, we considered the MMD(·) divergence function [75].

The ELBO term proposed in this work, which is the measure maximised during the training of AEs, is:

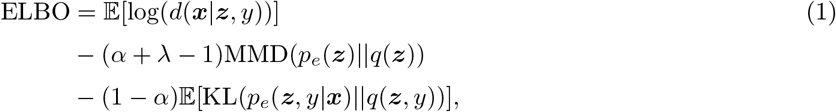

where KL(*·*) is the Kullback-Leibler divergence [76] between two distributions. All the mathematical details required to derive the generalised formula shown in Equation 1 can be found in the Additional file 1.

Equation 1 allows the user to easily exploit VAE, MMDAE, MMDVAE, GMVAE, GMMMD, and GMMMMDVAE (see Table 1).

**Table 1.** Setting of *α, λ*, and *K* to obtain the desired AE.

| | $\alpha$ | $\lambda$ | $K$ |
| --- | --- | --- | --- |
| VAE | 0 | 1 | 1 |
| MMDAE | 1 | 1 | 1 |
| MMDVAE | 0 | 2 | 1 |
| GMVAE | 0 | 1 | $> 1$ |
| GMMMD | 1 | 1 | $> 1$ |
| GMMMDVAE | 0 | 2 | $> 1$ |

### Availability and requirements

scAEspy is written in Python programming language (v.3.6.5) and it relies on TensorFlow (v.1.12.0), an open-source and massively used machine learning library [33]. scAEspy requires the following Python libraries: NumPy, SciKit-Learn, Matplotlib, and Seaborn. scAEspy’s open-source code is available on GitLab: https://gitlab.com/cvejic-group/scaespy under the GPL-3 license. scAEspy can be easily installed using the Python package installer pip, which allows the user to use scAEspy either as a standalone tool exploiting the command line interface, or as a Python package that can be integrated into Python scripts and Jupyter Notebooks.

The repository contains all the scripts, code and Jupyter Notebooks used to obtain the results shown in the paper. In the provided Jupyter Notebooks, we show how easy it is to integrate scAEspy and Scanpy, and how the data can be visualised and explored by using both scAEspy and Scanpy’s functions. We also provide a detailed description of scAEspy’s parameters so that it can be used by both novice and expert researchers for downstream analyses.

## Declarations

### Ethics approval, accordance and consent to participate

Not applicable

## Consent for publication

Not applicable

## Availability of data and materials

The datasets analysed during the current study as well as all the developed code are available in the scAEspy repository: https://gitlab.com/cvejic-group/scaespy.

## Author’s contributions

AT conceived the project. AT and FR developed the software. AC, DB, and PL supervised the project and helped to interpret and present the results. AT performed all the tests and analysed the results. AT wrote the manuscript. AC, DB, and PL edited the manuscript. All authors read and approved the final manuscript.

## Acknowledgements

We thank Dr. Leonardo Rundo (Department of Radiology, University of Cambridge) for his critical comments.

## References

1. Gladka, M.M., Molenaar, B., De Ruiter, H., Van Der Elst, S., Tsui, H., Versteeg, D., Lacraz, G.P., Huibers, M.M., Van Oudenaarden, A., Van Rooij, E.: Single-cell sequencing of the healthy and diseased heart reveals cytoskeleton-associated protein 4 as a new modulator of fibroblasts activation. Circulation 138(2), 166–180 (2018). doi:10.1161/CIRCULATIONAHA.117.030742

2. Keren-Shaul, H., Spinrad, A., Weiner, A., Matcovitch-Natan, O., Dvir-Szternfeld, R., Ulland, T.K., David, E., Baruch, K., Lara-Astaiso, D., Toth, B., et al.: A unique microglia type associated with restricting development of Alzheimer’s disease. Cell 169(7), 1276–1290 (2017). doi:10.1016/j.cell.2017.05.018

3. Lähnemann, D., Köster, J., Szczurek, E., McCarthy, D.J., Hicks, S.C., Robinson, M.D., Vallejos, C.A., Campbell, K.R., Beerenwinkel, N., Mahfouz, A., et al.: Eleven grand challenges in single-cell data science. Genome Biol. 21(1), 1–35 (2020). doi:10.1186/s13059-020-1926-6

4. Steinbach, M., Ertöz, L., Kumar, V.: The challenges of clustering high dimensional data. In: New Directions in Statistical Physics: Econophysics, Bioinformatics, and Pattern Recognition, pp. 273–309. Springer, Berlin, Heidelberg (2004). doi:10.1007/978-3-662-08968-2_16

5. Wold, S., Esbensen, K., Geladi, P.: Principal component analysis. Chemom Intell Lab Syst. 2(1-3), 37–52 (1987). doi:10.1016/0169-7439(87)80084-9

6. Maaten, L.v.d., Hinton, G.: Visualizing data using t-SNE. J Mach Learn Res. 9(Nov), 2579–2605 (2008)

7. Becht, E., McInnes, L., Healy, J., Dutertre, C.-A., Kwok, I.W., Ng, L.G., Ginhoux, F., Newell, E.W.: Dimensionality reduction for visualizing single-cell data using UMAP. Nat Biotechnol. 37(1), 38 (2019). doi:10.1038/nbt.4314

8. Luecken, M.D., Theis, F.J.: Current best practices in single-cell RNA-seq analysis: a tutorial. Mol Syst Biol. 15(6), 8746 (2019). doi:10.15252/msb.20188746

9. Hwang, B., Lee, J.H., Bang, D.: Single-cell RNA sequencing technologies and bioinformatics pipelines. Exp Mol Med. 50(8), 96 (2018). doi:10.1038/s12276-018-0071-8

10. Luecken, M.D., Buttner, M., Chaichoompu, K., Danese, A., Interlandi, M., Müller, M.F., Strobl, D.C., Zappia, L., Dugas, M., Colomé-Tatché, M., et al.: Benchmarking atlas-level data integration in single-cell genomics. BioRxiv (2020). doi:10.1101/2020.05.22.111161

11. Tran, H.T.N., Ang, K.S., Chevrier, M., Zhang, X., Lee, N.Y.S., Goh, M., Chen, J.: A benchmark of batch-effect correction methods for single-cell RNA sequencing data. Genome Biol. 21(1), 1–32 (2020). doi:10.1186/s13059-019-1850-9

12. Leek, J.T., Scharpf, R.B., Bravo, H.C., Simcha, D., Langmead, B., Johnson, W.E., Geman, D., Baggerly, K., Irizarry, R.A.: Tackling the widespread and critical impact of batch effects in high-throughput data. Nat Rev Genet. 11(10), 733–739 (2010). doi:10.1038/nrg2825

13. Butler, A., Hoffman, P., Smibert, P., Papalexi, E., Satija, R.: Integrating single-cell transcriptomic data across different conditions, technologies, and species. Nat Biotechnol. 36(5), 411 (2018). doi:10.1038/nbt.4096

14. Bacher, R., Kendziorski, C.: Design and computational analysis of single-cell RNA-sequencing experiments. Genome Biol. 17(1), 63 (2016). doi:10.1186/s13059-016-0927-y

15. Ding, J., Condon, A., Shah, S.P.: Interpretable dimensionality reduction of single cell transcriptome data with deep generative models. Nat Commun. 9(1), 2002 (2018). doi:10.1038/s41467-018-04368-5

16. Eraslan, G., Simon, L.M., Mircea, M., Mueller, N.S., Theis, F.J.: Single-cell RNA-seq denoising using a deep count autoencoder. Nat Commun. 10(1), 390 (2019). doi:10.1038/s41467-018-07931-2

17. Kingma, D.P., Welling, M.: Auto-encoding variational bayes. arXiv preprint arXiv:1312.6114 (2013)

18. Lopez, R., Regier, J., Cole, M.B., Jordan, M.I., Yosef, N.: Deep generative modeling for single-cell transcriptomics. Nat Methods 15(12), 1053 (2018). doi:10.1038/s41592-018-0229-2

19. Svensson, V., Gayoso, A., Yosef, N., Pachter, L.: Interpretable factor models of single-cell RNA-seq via variational autoencoders. Bioinformatics 36(11), 3418–3421 (2020). doi:10.1093/bioinformatics/btaa169

20. Grønbech, C.H., Vording, M.F., Timshel, P.N., Sønderby, C.K., Pers, T.H., Winther, O.: scVAE: Variational auto-encoders for single-cell gene expression data. Bioinformatics (2020). doi:10.1093/bioinformatics/btaa293

21. Tran, D., Nguyen, H., Tran, B., La Vecchia, C., Luu, H.N., Nguyen, T.: Fast and precise single-cell data analysis using a hierarchical autoencoder. Nat Commun. 12(1), 1–10 (2021). doi:10.1038/s41467-021-21312-2

22. Rousseeuw, J.: A graphical aid to the interpretation and validation of cluster analysis. J Comput Appl Math. 20, 53–65 (1989)

23. Bica, I., Andrés-Terré, H., Cvejic, A., Liò, P.: Unsupervised generative and graph representation learning for modelling cell differentiation. Sci Rep. 10(1), 1–13 (2020). doi:10.1038/s41598-020-66166-8

24. Wang, D., Gu, J.: VASC: dimension reduction and visualization of single-cell RNA-seq data by deep variational autoencoder. Genom Proteom Bioinf. 16(5), 320–331 (2018). doi:10.1016/j.gpb.2018.08.003

25. Lin, E., Mukherjee, S., Kannan, S.: A deep adversarial variational autoencoder model for dimensionality reduction in single-cell RNA sequencing analysis. BMC bioinformatics 21(1), 1–11 (2020). doi:10.1186/s12859-020-3401-5

26. Geddes, T.A., Kim, T., Nan, L., Burchfield, J.G., Yang, J.Y., Tao, D., Yang, P.: Autoencoder-based cluster ensembles for single-cell RNA-seq data analysis. BMC bioinformatics 20(19), 660 (2019). doi:10.1186/s12859-019-3179-5

27. Talwar, D., Mongia, A., Sengupta, D., Majumdar, A.: AutoImpute: Autoencoder based imputation of single-cell RNA-seq data. Sci Rep. 8(1), 16329 (2018). doi:10.1038/s41598-018-34688-x

28. Sun, S., Liu, Y., Shang, X.: Deep generative autoencoder for low-dimensional embedding extraction from single-cell RNAseq data. In: Proceedings of the IEEE International Conference on Bioinformatics and Biomedicine, pp. 1365–1372 (2019). doi:10.1109/BIBM47256.2019.8983289. IEEE

29. Badsha, M.B., Li, R., Liu, B., Li, Y.I., Xian, M., Banovich, N.E., Fu, A.Q.: Imputation of single-cell gene expression with an autoencoder neural network. Quant Biol., 1–17 (2020). doi:10.1007/s40484-019-0192-7

30. Rao, J., Zhou, X., Lu, Y., Zhao, H., Yang, Y.: Imputing single-cell RNA-seq data by combining graph convolution and autoencoder neural networks. BioRxiv (2020). doi:10.1101/2020.02.05.935296

31. Wolf, F.A., Angerer, P., Theis, F.J.: SCANPY: large-scale single-cell gene expression data analysis. Genome Biol. 19(1), 15 (2018). doi:10.1186/s13059-017-1382-0

32. Satija, R., Farrell, J.A., Gennert, D., Schier, A.F., Regev, A.: Spatial reconstruction of single-cell gene expression data. Nat Biotechnol. 33(5), 495 (2015). doi:10.1038/nbt.3192

33. Abadi, M., Barham, P., Chen, J., Chen, Z., Davis, A., Dean, J., Devin, M., Ghemawat, S., Irving, G., Isard, M., et al.: Tensorflow: A system for large-scale machine learning. In: Proceedings of the Symposium on Operating Systems Design and Implementation), pp. 265–283 (2016)

34. Paszke, A., Gross, S., Chintala, S., Chanan, G., Yang, E., DeVito, Z., Lin, Z., Desmaison, A., Antiga, L., Lerer, A.: Automatic differentiation in PyTorch. In: Proceedings of the Conference on Advances in Neural Information Processing Systems (2017)

35. Zhao, S., Song, J., Ermon, S.: Infovae: Balancing learning and inference in variational autoencoders. In: Proceedings of the AAAI Conference on Artificial Intelligence, vol. 33, pp. 5885–5892 (2019)

36. Polański, K., Young, M.D., Miao, Z., Meyer, K.B., Teichmann, S.A., Park, J.-E.: BBKNN: fast batch alignment of single cell transcriptomes. Bioinformatics 36(3), 964–965 (2020). doi:10.1093/bioinformatics/btz625

37. Korsunsky, I., Millard, N., Fan, J., Slowikowski, K., Zhang, F., Wei, K., Baglaenko, Y., Brenner, M., Loh, P.-r., Raychaudhuri, S.: Fast, sensitive and accurate integration of single-cell data with Harmony. Nat Methods 16, 1289–1296 (2019). doi:10.1038/s41592-019-0619-0

38. Johnson, W.E., Li, C., Rabinovic, A.: Adjusting batch effects in microarray expression data using empirical Bayes methods. Biostatistics 8(1), 118–127 (2007). doi:10.1093/biostatistics/kxj037

39. Leek, J.T., Johnson, W.E., Parker, H.S., Jaffe, A.E., Storey, J.D.: The sva package for removing batch effects and other unwanted variation in high-throughput experiments. Bioinformatics 28(6), 882–883 (2012). doi:10.1093/bioinformatics/bts034

40. Pedersen, B.: Python implementation of ComBat. GitHub (2012)

41. Kang, H.M., Subramaniam, M., Targ, S., Nguyen, M., Maliskova, L., McCarthy, E., Wan, E., Wong, S., Byrnes, L., Lanata, C.M., et al.: Multiplexed droplet single-cell RNA-sequencing using natural genetic variation. Nat Biotechnol. 36(1), 89 (2018). doi:10.1038/nbt.4042

42. Grün, D., Muraro, M.J., Boisset, J.-C., Wiebrands, K., Lyubimova, A., Dharmadhikari, G., van den Born, M., van Es, J., Jansen, E., Clevers, H., et al.: De novo prediction of stem cell identity using single-cell transcriptome data. Cell Stem Cell 19(2), 266–277 (2016). doi:10.1016/j.stem.2016.05.010

43. Muraro, M.J., Dharmadhikari, G., Grün, D., Groen, N., Dielen, T., Jansen, E., van Gurp, L., Engelse, M.A., Carlotti, F., de Koning, E.J., et al.: A single-cell transcriptome atlas of the human pancreas. Cell Syst. 3(4), 385–394 (2016). doi:10.1016/j.cels.2016.09.002

44. Lawlor, N., George, J., Bolisetty, M., Kursawe, R., Sun, L., Sivakamasundari, V., Kycia, I., Robson, P., Stitzel, M.L.: Single-cell transcriptomes identify human islet cell signatures and reveal cell-type–specific expression changes in type 2 diabetes. Genome Res. 27(2), 208–222 (2017). doi:10.1101/gr.212720.116

45. Segerstolpe, Å., Palasantza, A., Eliasson, P., Andersson, E.-M., Andréasson, A.-C., Sun, X., Picelli, S., Sabirsh, A., Clausen, M., Bjursell, M.K., et al.: Single-cell transcriptome profiling of human pancreatic islets in health and type 2 diabetes. Cell Metab. 24(4), 593–607 (2016). doi:10.1016/j.cmet.2016.08.020

46. Stuart, T., Butler, A., Hoffman, P., Hafemeister, C., Papalexi, E., Mauck III W.M., Hao, Y., Stoeckius, M., Smibert, P., Satija, R.: Comprehensive integration of single-cell data. Cell (2019). doi:10.1016/j.cell.2019.05.031

47. Han, X., Wang, R., Zhou, Y., Fei, L., Sun, H., Lai, S., Saadatpour, A., Zhou, Z., Chen, H., Ye, F., et al.: Mapping the mouse cell atlas by microwell-seq. Cell 172(5), 1091–1107 (2018). doi:10.1016/j.cell.2018.02.001

48. Consortium, T.M., et al.: Single-cell transcriptomics of 20 mouse organs creates a Tabula Muris. Nature 562, 367–372 (2018). doi:10.1038/s41586-018-0590-4

49. Hubert, L., Arabie, P.: Comparing partitions. J Classif. 2(1), 193–218 (1985). doi:10.1007/BF01908075

50. Strehl, A., Ghosh, J.: Cluster ensembles—a knowledge reuse framework for combining multiple partitions. J Mach Learn Res. 3(Dec), 583–617 (2002)

51. Fowlkes, E.B., Mallows, C.L.: A method for comparing two hierarchical clusterings. J Am Stat Assoc. 78(383), 553–569 (1983)

52. Vinh, N.X., Epps, J., Bailey, J.: Information theoretic measures for clusterings comparison: Variants, properties, normalization and correction for chance. J Mach Learn Res. 11(Oct), 2837–2854 (2010)

53. Rosenberg, A., Hirschberg, J.: V-measure: A conditional entropy-based external cluster evaluation measure. In: Proceedings of the Conference on Empirical Methods in Natural Language Processing and Computational Natural Language Learning, pp. 410–420 (2007)

54. Macosko, E.Z., Basu, A., Satija, R., Nemesh, J., Shekhar, K., Goldman, M., Tirosh, I., Bialas, A.R., Kamitaki, N., Martersteck, E.M., et al.: Highly parallel genome-wide expression profiling of individual cells using nanoliter droplets. Cell 161(5), 1202–1214 (2015). doi:10.1016/j.cell.2015.05.002

55. Klein, A.M., Mazutis, L., Akartuna, I., Tallapragada, N., Veres, A., Li, V., Peshkin, L., Weitz, D.A., Kirschner, M.W.: Droplet barcoding for single-cell transcriptomics applied to embryonic stem cells. Cell 161(5), 1187–1201 (2015). doi:10.1016/j.cell.2015.04.044

56. Hashimshony, T., Wagner, F., Sher, N., Yanai, I.: CEL-Seq: single-cell RNA-Seq by multiplexed linear amplification. Cell Rep. 2(3), 666–673 (2012). doi:10.1016/j.celrep.2012.08.003

57. Hashimshony, T., Senderovich, N., Avital, G., Klochendler, A., de Leeuw, Y., Anavy, L., Gennert, D., Li, S., Livak, K.J., Rozenblatt-Rosen, O., et al.: CEL-Seq2: sensitive highly-multiplexed single-cell RNA-Seq. Genome Biol. 17(1), 77 (2016). doi:10.1186/s13059-016-0938-8

58. Zheng, G.X., Terry, J.M., Belgrader, P., Ryvkin, P., Bent, Z.W., Wilson, R., Ziraldo, S.B., Wheeler, T.D., McDermott, G.P., Zhu, J., et al.: Massively parallel digital transcriptional profiling of single cells. Nat Commun. 8, 14049 (2017). doi:10.1038/ncomms14049

59. Gierahn, T.M., Wadsworth II M.H., Hughes, T.K., Bryson, B.D., Butler, A., Satija, R., Fortune, S., Love, J.C., Shalek, A.K.: Seq-Well: portable, low-cost RNA sequencing of single cells at high throughput. Nat Methods 14(4), 395 (2017). doi:10.1038/nmeth.4179

60. Islam, S., Kjällquist, U., Moliner, A., Zajac, P., Fan, J.-B., Lönnerberg, P., Linnarsson, S.: Characterization of the single-cell transcriptional landscape by highly multiplex RNA-seq. Genome Res. 21(7), 1160–1167 (2011). doi:10.1101/gr.110882.110

61. Ramsköld, D., Luo, S., Wang, Y.-C., Li, R., Deng, Q., Faridani, O.R., Daniels, G.A., Khrebtukova, I., Loring, J.F., Laurent, L.C., et al.: Full-length mRNA-Seq from single-cell levels of RNA and individual circulating tumor cells. Nat Biotechnol. 30(8), 777 (2012). doi:10.1038/nbt.2282

62. Picelli, S., Faridani, O.R., Björklund, Å.K., Winberg, G., Sagasser, S., Sandberg, R.: Full-length RNA-seq from single cells using smart-seq2. Nat Protoc. 9(1), 171 (2014). doi:10.1038/nprot.2014.006

63. Jaitin, D.A., Kenigsberg, E., Keren-Shaul, H., Elefant, N., Paul, F., Zaretsky, I., Mildner, A., Cohen, N., Jung, S., Tanay, A., et al.: Massively parallel single-cell RNA-seq for marker-free decomposition of tissues into cell types. Science 343(6172), 776–779 (2014). doi:10.1126/science.1247651

64. Traag, V.A., Waltman, L., van Eck, N.J.: From Louvain to Leiden: guaranteeing well-connected communities. Sci Rep. 9 (2019). doi:10.1038/s41598-019-41695-z

65. Mann, H.B., Whitney, D.R.: On a test of whether one of two random variables is stochastically larger than the other. Ann Math Stat., 50–60 (1947)

66. Wilcoxon, F.: Individual comparisons by ranking methods. In: Breakthroughs in Statistics, pp. 196–202. Springer, New York, NY (1992). doi:10.1007/978-1-4612-4380-9_16

67. Dunn, O.J.: Multiple comparisons among means. J Am Stat Assoc. 56(293), 52–64 (1961)

68. Ma, F., Pellegrini, M.: ACTINN: Automated identification of cell types in single cell RNA sequencing. Bioinformatics (2019). doi:10.1093/bioinformatics/btz592

69. Ranzoni, A.M., Tangherloni, A., Berest, I., Riva, S.G., Myers, B., Strzelecka, P.M., Xu, J., Panada, E., Mohorianu, I., Zaugg, J.B., et al.: Integrative single-cell rna-seq and atac-seq analysis of human developmental hematopoiesis. Cell Stem Cell 28(3), 472–487 (2021). doi:10.1016/j.stem.2020.11.015

70. Simidjievski, N., Bodnar, C., Tariq, I., Scherer, P., Andres Terre, H., Shams, Z., Jamnik, M., Liò, P.: Variational autoencoders for cancer data integration: design principles and computational practice. Front Genet. 10, 1205 (2019). doi:10.3389/fgene.2019.01205

71. Trębacz, M., Shams, Z., Jamnik, M., Scherer, P., Simidjievski, N., Terre, H.A., Liò, P.: Using ontology embeddings for structural inductive bias in gene expression data analysis. arXiv preprint arXiv:2011.10998 (2020)

72. Ashburner, M., Ball, C.A., Blake, J.A., Botstein, D., Butler, H., Cherry, J.M., Davis, A.P., Dolinski, K., Dwight, S.S., Eppig, J.T., et al.: Gene ontology: tool for the unification of biology. Nat Genet. 25(1), 25–29 (2000)

73. Raudvere, U., Kolberg, L., Kuzmin, I., Arak, T., Adler, P., Peterson, H., Vilo, J.: g:Profiler: a web server for functional enrichment analysis and conversions of gene lists (2019 update). Nucl Acids Res. 47(W1), 191–198 (2019). doi:10.1093/nar/gkz369

74. Chen, X., Miragaia, R.J., Natarajan, K.N., Teichmann, S.A.: A rapid and robust method for single cell chromatin accessibility profiling. Nat Commun. 9(1), 5345 (2018). doi:10.1038/s41467-018-07771-0

75. Gretton, A., Borgwardt, K., Rasch, M., Schölkopf, B., Smola, A.J.: A kernel method for the two-sample-problem. In: Proceedings of the Conference on Advances in Neural Information Processing Systems, pp. 513–520 (2007)

76. Kullback, S., Leibler, R.A.: On information and sufficiency. Ann Math Statist. 22(1), 79–86 (1951). doi:10.1214/aoms/1177729694

77. Risso, D., Perraudeau, F., Gribkova, S., Dudoit, S., Vert, J.-P.: A general and flexible method for signal extraction from single-cell RNA-seq data. Nat Commun. 9(1), 1–17 (2018). doi:10.1038/s41467-017-02554-5

78. Hie, B., Bryson, B., Berger, B.: Efficient integration of heterogeneous single-cell transcriptomes using Scanorama. Nat Biotechnol. 37(6), 685–691 (2019). doi:10.1038/s41587-019-0113-3

79. Haghverdi, L., Lun, A.T., Morgan, M.D., Marioni, J.C.: Batch effects in single-cell RNA-sequencing data are corrected by matching mutual nearest neighbors. Nat Biotechnol. 36(5), 421 (2018). doi:10.1038/nbt.4091

80. Kingma, D.P., Ba, J.: Adam: A method for stochastic optimization. arXiv preprint arXiv:1412.6980 (2014)

