## Additional file 1 for "Analysis of single-cell RNA sequencing data based on autoencoders"

#### Mathematical formulation of the proposed autoencoders

In what follows, we use the notation proposed in [1] to derive the extension of the Mean Maximum Discrepancy Autoencoder (MMDAE) and Mean Maximum Discrepancy Variational Autoencoder (MMDVAE) [1] with multiple Gaussian distributions.

$p^*(\mathbf{x})$ is the unknown probability in the input space over which the optimisation problem is formulated. A similarity measure $\mathcal{L}$ between the input and the output of the autoencoders (AEs) with respect to the distribution $p^*(\mathbf{x})$ is maximised:

$$\arg\max_{\boldsymbol{\phi},\boldsymbol{\theta}} \mathbb{E}[\mathcal{L}(\boldsymbol{x}, d_{\boldsymbol{\theta}}(e_{\boldsymbol{\phi}}(\boldsymbol{x})))],$$

where $\boldsymbol{\phi}$ and $\boldsymbol{\theta}$ are the weights of the encoder and decoder networks, respectively; $e_{\boldsymbol{\phi}} : \boldsymbol{x} \mapsto \boldsymbol{z}$ and $d_{\boldsymbol{\theta}} : \boldsymbol{z} \mapsto \boldsymbol{x}$, where $\boldsymbol{z}$ is the latent representation of $\boldsymbol{x}$ and $|\boldsymbol{z}| \leq |\boldsymbol{x}|$.

In variational AEs, the input $\boldsymbol{x}$ is mapped into a probability distribution over the latent space. $e(\boldsymbol{z}|\boldsymbol{x})$ defines a distribution over the latent space that depends on the input $\boldsymbol{x}$ drawn from $p^*(\mathbf{x})$. Altogether, $p^*(\mathbf{x})$ and $e(\boldsymbol{z}|\boldsymbol{x})$ define the joint distribution $p_e(\boldsymbol{x}, \boldsymbol{z}) = e(\boldsymbol{z}|\boldsymbol{x})p^*(\mathbf{x})$, whose marginal and conditional distributions are defined as:

$$p_e(\boldsymbol{z}) = \int p_e(\boldsymbol{x}, \boldsymbol{z})d\boldsymbol{x} = \int p_e(\boldsymbol{z}|\boldsymbol{x})p^*(\boldsymbol{x})d\boldsymbol{x}$$

$$p_e(\boldsymbol{x}|\boldsymbol{z}) = \frac{p_e(\boldsymbol{x}, \boldsymbol{z})}{p_e(\boldsymbol{z})}.$$

Since the representation $\boldsymbol{z}$ of $\boldsymbol{x}$ should maintain as much as possible the "amount of information" held in $\boldsymbol{x}$, the mutual information $I(\boldsymbol{x}; \boldsymbol{z})$ can be used to measure the representation $\boldsymbol{z}$ of $\boldsymbol{x}$. Specifically, for any distribution $q(\boldsymbol{z})$ in the latent space, the mutual information between $p_e(\boldsymbol{z})$ and $p^*(\boldsymbol{x})$ can be bounded below as:

$$I(\boldsymbol{x}; \boldsymbol{z}) = \mathrm{KL}(p_e(\boldsymbol{x}, \boldsymbol{z})||p^*(\boldsymbol{x})p_e(\boldsymbol{z})) \leq \mathbb{E}[\mathrm{KL}(e(\boldsymbol{z}|\boldsymbol{x})||q(\boldsymbol{z}))],$$

where $\mathrm{KL}(\cdot)$ is the Kullback–Leibler divergence [2] between two distributions. $I(\boldsymbol{x}; \boldsymbol{z})$ can be also bounded above, for any conditional distribution $d(\boldsymbol{x}|\boldsymbol{z})$, as:

$$I(\boldsymbol{x}; \boldsymbol{z}) = \mathrm{KL}(p_e(\boldsymbol{x}, \boldsymbol{z})||p^*(\boldsymbol{x})p_e(\boldsymbol{z})) \geq \mathbb{E}\left[\log\left(\frac{d(\boldsymbol{x}|\boldsymbol{z})}{p^*(\boldsymbol{x})}\right)\right].$$

Combining the provided definitions, we obtain that:

$$\mathbb{E}\left[\log\left(\frac{d(\boldsymbol{x}|\boldsymbol{z})}{p^*(\boldsymbol{x})}\right)\right] \leq I(\boldsymbol{x};\boldsymbol{z}) \leq \mathbb{E}[\mathrm{KL}(e(\boldsymbol{z}|\boldsymbol{x})||q(\boldsymbol{z}))].$$

The lower bound can be further decomposed by means of algebraic manipulations as

$$\mathbb{E}\left[\log\left(\frac{d(\boldsymbol{x}|\boldsymbol{z})}{p^*(\boldsymbol{x})}\right)\right] = \mathbb{E}[\log(d(\boldsymbol{x}|\boldsymbol{z})) + H(p^*(\boldsymbol{x}))],$$

where $H(p^*(\boldsymbol{x}))]$ is the entropy of $p^*(\boldsymbol{x})$. By following the definition provided in [1], the ELBO term, which is the measure maximised during the training of VAE, can be written as:

$$\mathrm{ELBO} = -\mathrm{KL}(p_e(\mathbf{z})||q(\mathbf{z})) - H(p^*(\boldsymbol{x})) - \mathbb{E}[\mathrm{KL}(p_e(\boldsymbol{x}|\boldsymbol{z})||d(\boldsymbol{x}|\boldsymbol{z}))].$$

In MMDVAE [1], $\mathrm{KL}(p_e(\mathbf{z})||q(\mathbf{z}))$ is multiplied by a positive factor $\lambda$ and $I(\boldsymbol{x};\boldsymbol{z})$, weighted by a positive factor $\alpha$, is added to the ELBO term, obtaining:

$$\begin{aligned}\mathrm{ELBO} = &-\lambda\mathrm{KL}(p_e(\boldsymbol{z})||q(\boldsymbol{z}))\\ &- H(p^*(\boldsymbol{x})\\ &- \mathbb{E}[\mathrm{KL}(p_e(\boldsymbol{x}|\boldsymbol{z})||d(\boldsymbol{x}|\boldsymbol{z}))]\\ &+ \alpha I(\boldsymbol{x};\boldsymbol{z}).\end{aligned}$$

By applying algebraic manipulations, the ELBO term of MMDVAE can be written as:

$$\begin{aligned}\mathrm{ELBO} = &\,\mathbb{E}[\log(d(\boldsymbol{x}|\boldsymbol{z}))]\\ &- (\alpha+\lambda-1)\mathrm{KL}(p_e(\boldsymbol{z})||q(\boldsymbol{z}))\\ &- (1-\alpha)\mathbb{E}[\mathrm{KL}(p_e(\boldsymbol{z}|\boldsymbol{x})||q(\boldsymbol{z})).\end{aligned} \tag{1}$$

In MMDVAE, the term $\mathrm{KL}(p_e(\boldsymbol{z})||q(\boldsymbol{z}))$ is replaced with $\mathrm{DSD}(p_e(\boldsymbol{z})||q(\boldsymbol{z}))$, where $DSD(\cdot)$ is a general strict divergence function. $\mathrm{DSD}(p_e(\boldsymbol{z})||q(\boldsymbol{z})) = 0$ if and only if $p_e(\cdot) = q(\cdot)$. Notice that, the KL is a strict divergence function. MMDVAE exploits the Maximum Mean Discrepancy $\mathrm{MMD}(\cdot)$ divergence function [3]. A kernel trick is used to define the following divergence function between two distributions $p_e(\boldsymbol{z})$ and $q(\boldsymbol{z})$:

$$\begin{aligned}\mathrm{MMD}(p_e(\boldsymbol{z})||q(\boldsymbol{z})) = &\,\mathbb{E}_{p_e(\boldsymbol{z}),p(\boldsymbol{z'})}[\mathcal{K}(\boldsymbol{z},\boldsymbol{z'})]\\ &+ \mathbb{E}_{q(\boldsymbol{z}),q(\boldsymbol{z'})}[\mathcal{K}(\boldsymbol{z},\boldsymbol{z'})]\\ &- 2\mathbb{E}_{p_e(\boldsymbol{z}),q(\boldsymbol{z'})}[\mathcal{K}(\boldsymbol{z},\boldsymbol{z'})],\end{aligned}$$

where $\mathcal{K}(\boldsymbol{z},\boldsymbol{z'})$ can be any desired universal kernel. Here, we considered the Gaussian kernel

$$\mathcal{K}(\boldsymbol{z},\boldsymbol{z'}) = e^{-\frac{||\boldsymbol{z}-\boldsymbol{z'}||}{2\sigma^2}}.$$

We extended the ELBO term shown in Eq. 1 such that multiple Gaussian distributions can be used in the latent representation $\boldsymbol{z}$. In addition, we introduced a learnable mixture distribution for $q(\boldsymbol{z})$, whereas $p_e(\boldsymbol{z}|\boldsymbol{x})$ is defined to be a learnable mixture distribution with the same number of components.

In GMVAE [4], the encoder function outputs the following two conditional distributions $e(\boldsymbol{z}, y|\boldsymbol{x})$ and $e(\boldsymbol{z}|\boldsymbol{x}, y)$, where $y \in \{1, \ldots, K\}$ is a categorical random variable and $K$ corresponds to the number of desired Gaussian distributions. We obtain that

$$\begin{aligned} p_e(\boldsymbol{z}, y|\boldsymbol{x}) &= \frac{p_e(\boldsymbol{z}, y, \boldsymbol{x})}{p^*(\boldsymbol{x})} \\ &= e(\boldsymbol{z}|y, \boldsymbol{x})e(y|\boldsymbol{x}). \end{aligned}$$

Since $p_e(\boldsymbol{z}, y|\boldsymbol{x})$ is fully determined by the output distributions of the encoder, we can refer to $p_e(\boldsymbol{z}, y|\boldsymbol{x})$ with $e(\boldsymbol{z}, y|\boldsymbol{x})$.

Modelling $e(y|\boldsymbol{x})$ as a categorical distribution that can assume values in $\{1, \ldots, K\}$, and $e(\boldsymbol{z}|\boldsymbol{x}, y)$ as a diagonal Gaussian distribution for each possible value assumed by $y$, the marginal conditional distribution $p_e(\boldsymbol{z}|\boldsymbol{x})$ is a Gaussian mixture distribution of $K$ components, namely:

$$p_e(\boldsymbol{z}|\boldsymbol{x}) = \sum_{y=1}^{K} e(\boldsymbol{z}|y, \boldsymbol{x})e(y|\boldsymbol{x}).$$

Similarly, $q(\boldsymbol{z})$ is modelled as a Gaussian mixture distribution by using another variable $y \in \{1, \ldots, K\}$, with a categorical distribution $q(y)$, and considering the conditional distribution $q(\boldsymbol{z}|y)$ as a diagonal Gaussian distribution for each possible value of $y$. The ELBO term of GMVAE is:

$$ELBO = \mathbb{E}\left[\mathbb{E}\left[d(\boldsymbol{x}|y, \boldsymbol{z}) - \log\left(\frac{e(\boldsymbol{z}, y|\boldsymbol{x})}{q(\boldsymbol{z}, y)}\right)\right]\right],$$

and it can be rewritten by using the notation proposed in [1] and by algebraic manipulations as:

$$\begin{aligned} ELBO &= -\mathrm{KL}(p_e(\boldsymbol{z}, y)||q(\boldsymbol{z}, y)) \\ &= -\mathbb{E}[\mathrm{KL}(p_e(\boldsymbol{x}|\boldsymbol{z}, y)||d(\boldsymbol{x}|\boldsymbol{z}, y))] \\ &= -H(p^*(\boldsymbol{x})). \end{aligned}$$

Starting from this definition, we can add the mutual information $I(\boldsymbol{x}; (y, \boldsymbol{z}))$ term, weighted by a positive scalar factor $\alpha$, and $\mathrm{KL}(p_e(\boldsymbol{z}, y)||q(\boldsymbol{z}, y))$ is weighted by a positive factor $\lambda$, obtaining:

$$\begin{aligned} \mathrm{ELBO} = &-\lambda \mathrm{KL}(p_e(\boldsymbol{z}, y)||q(\boldsymbol{z}, y)) \\ &- H(p^*(\boldsymbol{x})) \\ &- \mathbb{E}[\mathrm{KL}(p_e(\boldsymbol{x}|\boldsymbol{z}, y)||d(\boldsymbol{x}|\boldsymbol{z}, y))] \\ &+ \alpha I(\boldsymbol{x}; (y, \boldsymbol{z})), \end{aligned}$$

where

$$I(\boldsymbol{x}; (y, \boldsymbol{z})) = \mathbb{E}\left[\log \frac{p_e(\boldsymbol{x}, y, \boldsymbol{z})}{p^*(\boldsymbol{x})p_e(\boldsymbol{z}, y))}\right].$$

By applying algebraic manipulations, we can rewrite the ELBO term as:

$$\begin{aligned} \mathrm{ELBO} = &\mathbb{E}[\log(d(\boldsymbol{x}|\boldsymbol{z}, y))] \\ &- (\alpha + \lambda - 1)\mathrm{KL}(p_e(\boldsymbol{z}, y)||q(\boldsymbol{z}, y)) \\ &- (1 - \alpha)\mathbb{E}[\mathrm{KL}(p_e(\boldsymbol{z}, y|\boldsymbol{x})||q(\boldsymbol{z}, y))]. \end{aligned}$$

$\mathrm{KL}(p_e(\boldsymbol{z}, y)||q(\boldsymbol{z}, y))$ can be further decomposed as:

$$\mathrm{KL}(p_e(\boldsymbol{z}, y)||q(\boldsymbol{z}, y)) = \mathbb{E}[\mathrm{KL}(p_e(y|\boldsymbol{z})||q(y|\boldsymbol{z}))] + \mathrm{KL}(p_e(\boldsymbol{z})||q(\boldsymbol{z})),$$

so that the ELBO can be written as:

$$\begin{aligned}\mathrm{ELBO} = {} & \mathbb{E}[\log(d(\boldsymbol{x}|\boldsymbol{z}, y))] \\ & - (\alpha + \lambda - 1)\mathrm{KL}(p_e(\boldsymbol{z})||q(\boldsymbol{z})) \\ & - (1 - \alpha)\mathbb{E}[\mathrm{KL}(p_e(\boldsymbol{z}, y|\boldsymbol{x})||q(\boldsymbol{z}, y))].\end{aligned}$$

As in MMDVAE (see Eq. 1), we can replace $\mathrm{KL}(p_e(\boldsymbol{z})||q(\boldsymbol{z}))$ with a general strict divergence function. We considered the $\mathrm{MMD}(\cdot)$ term, obtaining the a general formulation for all the five AEs:

$$\begin{aligned}\mathrm{ELBO} = {} & \mathbb{E}[\log(d(\boldsymbol{x}|\boldsymbol{z}, y))] \\ & - (\alpha + \lambda - 1)\mathrm{MMD}(p_e(\boldsymbol{z})||q(\boldsymbol{z})) \\ & - (1 - \alpha)\mathbb{E}[\mathrm{KL}(p_e(\boldsymbol{z}, y|\boldsymbol{x})||q(\boldsymbol{z}, y))].\end{aligned} \tag{2}$$

We modified the $\mathrm{MMD}(p_e(\boldsymbol{z})||q(\boldsymbol{z}))$ such that it is not necessary to sample from the Gaussian mixture distribution $e(\boldsymbol{z}|\boldsymbol{x})$ or from the posterior $q(\boldsymbol{z})$. Our modification allows for sampling from the single Gaussian distributions that form the mixtures. We used the the reparametrization trick proposed in [5] so that $\mathrm{MMD}(p_e(\boldsymbol{z})||q(\boldsymbol{z}))$ can be approximated. Specifically, $\mathbb{E}_{p_e(\boldsymbol{z}),p(\boldsymbol{z'})}[\mathcal{K}(\boldsymbol{z}, \boldsymbol{z'})]$ can be approximated as:

$$\frac{1}{N^2}\sum_{i=1}^{N}\sum_{j=1}^{N}\sum_{y=1}^{K}\sum_{y'=1}^{K} p_e(y, x_i)p_e(y', x'_j)\mathbb{E}_{p_e(\boldsymbol{z}|x_i,y),p_e(\boldsymbol{z'}|x'_j,y')}[\mathcal{K}(\boldsymbol{z}, \boldsymbol{z'})],$$

where $N$ is the number of provided samples (i.e., cells). The approximation of $\mathbb{E}_{q(\boldsymbol{z}),q(\boldsymbol{z'})}[\mathcal{K}(\boldsymbol{z}, \boldsymbol{z'})]$ is:

$$\sum_{y=1}^{K}\sum_{y'=1}^{K} q(y)q(y')\mathbb{E}_{q(\boldsymbol{z}|y),q(\boldsymbol{z'}|y')}[\mathcal{K}(\boldsymbol{z}, \boldsymbol{z'})].$$

Finally, $\mathbb{E}_{p_e(\boldsymbol{z}),q(\boldsymbol{z'})}[\mathcal{K}(\boldsymbol{z}, \boldsymbol{z'})]$ is approximated as:

$$\sum_{i=1}^{N}\sum_{y=1}^{K}\sum_{y'=1}^{K} p_e(y|x_i)q(y')\mathbb{E}_{p_e(\boldsymbol{z}|y,x_i),q(\boldsymbol{z'}|y')}[\mathcal{K}(\boldsymbol{z}, \boldsymbol{z'})].$$

To calculate the ELBO function described in Eq. 2, $\mathbb{E}[\mathrm{KL}(p_e(\boldsymbol{z}, y|\boldsymbol{x})||q(\boldsymbol{z}, y))]$ must be computable. We can rewrite it as follows:

$$\begin{aligned}\mathbb{E}[\mathrm{KL}(p_e(\boldsymbol{z}, y|\boldsymbol{x})||q(\boldsymbol{z}, y))] &= \mathbb{E}\left[\log\frac{p_e(\boldsymbol{z}|y, \boldsymbol{x})p_e(y|\boldsymbol{x})}{q(\boldsymbol{z}|y)q(y)}\right] \\ &= \mathbb{E}\left[\log\frac{p_e(\boldsymbol{z}|y, \boldsymbol{x})}{q(\boldsymbol{z}|y)}\right] + \mathbb{E}\left[\mathbb{E}\left[\log\frac{p_e(y|\boldsymbol{x})}{q(y)}\right]\right] \\ &= \mathbb{E}[\mathrm{KL}(p_e(y|\boldsymbol{x})||q(y))] + \mathbb{E}[\mathbb{E}[\mathrm{KL}(p_e(\boldsymbol{z}|\boldsymbol{x}, y)||q(\boldsymbol{z}|y))]].\end{aligned}$$

Considering the weights of $q(\boldsymbol{z})$ fixed to a uniform distribution, $\mathbb{E}[\mathrm{KL}(p_e(y|\boldsymbol{x})||q(y))]$ can be written as:

$$\mathbb{E}[\mathrm{KL}(p_e(y|\boldsymbol{x})||q(y))] = \mathbb{E}\left[\sum_{y=1}^{K} p_e(y|\boldsymbol{x})\log(p_e(y|\boldsymbol{x}))\right] + \log(K).$$

On the contrary, when the weights are learnable, $\mathbb{E}[\mathrm{KL}(p_e(y|\boldsymbol{x})||q(y))]$ can be analytically calculated as:

$$\mathbb{E}[\mathrm{KL}(p_e(y|\boldsymbol{x})||q(y))] = \mathbb{E}\left[\sum_{y=1}^{K} p_e(y|\boldsymbol{x})\log(p_e(y|\boldsymbol{x})) - \sum_{y=1}^{K} p_e(y|\boldsymbol{x})\log(q(y))\right].$$

Finally, $\mathrm{KL}(p_e(\boldsymbol{z}|\boldsymbol{x},y)||q(\boldsymbol{z}|y))$ can be calculated by the following approximation:

$$\mathrm{KL}(p_e(\boldsymbol{z}|\boldsymbol{x},y)||q(\boldsymbol{z}|y)) = \mathbb{E}\left[\log\frac{p_e(\boldsymbol{z}|\boldsymbol{x},y)}{q(\boldsymbol{z}|y)}\right] \approx \log\frac{p_e(\boldsymbol{z}|\boldsymbol{x},y)}{q(\boldsymbol{z}|y)}$$

### Additional Files

#### Additional file 2 — Excel file of the metrics calculated for the PBMC datasets

Each tab is related to a tested approach and shows the calculated metrics and used method.

#### Additional file 3 — Excel file of the metrics calculated for the PIC datasets

Each tab is related to a tested approach and shows the calculated metrics and used method.

#### Additional file 4 — Excel file of the metrics calculated for the MCA datasets

Each tab is related to a tested approach and shows the calculated metrics and used method.

### References


[1] Zhao, S., Song, J., Ermon, S.: Infovae: Balancing learning and inference in variational autoencoders. In: Proceedings of the AAAI Conference on Artificial Intelligence, vol. 33, pp. 5885–5892 (2019)

[2] Kullback, S., Leibler, R.A.: On information and sufficiency. Ann Math Statist. **22**(1), 79–86 (1951). doi:10.1214/aoms/1177729694

[3] Gretton, A., Borgwardt, K., Rasch, M., Schölkopf, B., Smola, A.J.: A kernel method for the two-sample-problem. In: Proceedings of the Conference on Advances in Neural Information Processing Systems, pp. 513–520 (2007)

[4] Grønbech, C.H., Vording, M.F., Timshel, P.N., Sønderby, C.K., Pers, T.H., Winther, O.: scVAE: Variational auto-encoders for single-cell gene expression data. Bioinformatics (2020). doi:10.1093/bioinformatics/btaa293

[5] Kingma, D.P., Welling, M.: Auto-encoding variational bayes. arXiv preprint arXiv:1312.6114 (2013)
